# Local, but not circulating, complement C3 governs immune checkpoint blockade efficacy

**DOI:** 10.1101/2025.05.12.653608

**Authors:** Yuki Miyai, Yukihiro Shiraki, Daisuke Sugiyama, Yoshitaka Sato, Katsuhiro Kato, Naoya Asai, Fuyang Cao, Nobuyoshi Nagao, Kana Tanabe, Masahiro Nakatochi, Tetsunari Hase, Toyofumi Fengshi Chen-Yoshikawa, Tomoko Kobayashi, Shintaro Iwama, Nobutoshi Esaki, Shinji Mii, Hiroshi Arima, Hiroshi Kimura, Masahide Takahashi, Yuichi Ando, Atsushi Enomoto

## Abstract

The roles of circulating complement C3, produced in hepatocytes, in immune defense are well defined in the literature. Interestingly, the C3 gene evolved before the circulatory system, suggesting an essential but yet unknown role of locally produced C3. Here, we investigated the effect of C3 on cancer immunotherapy and found that cancer-associated fibroblast (CAF)-derived C3, not hepatocyte-derived C3, determines the efficacy of immune checkpoint blockade (ICB). Tumors developed in CAF-specific C3 knockout mice exhibited resistance to anti-PD-1 therapy, with increased immunosuppressive M2-like macrophage infiltration. Mechanistically, the C3 degradation product iC3b suppressed myeloid cell infiltration via complement receptor 3 signaling, and the therapeutic targeting of this pathway restored ICB sensitivity of immunotherapy-resistant tumors. In human cancers, stromal C3 expression correlates with reduced M2-like macrophage infiltration and improved immunotherapeutic outcomes. These data demonstrate the role of locally produced C3 in the innate immune response, which may have been conserved across many organisms.

## INTRODUCTION

Innate immunity plays a crucial role in the body’s defense against pathogens^1,2^ and has recently emerged as a critical factor in the success of cancer immunotherapy^3^. Complement C3, a fundamental component of innate immunity, serves as the convergence point for all complement activation pathways, including the classical, alternative, and lectin pathways^1,2^. C3 has a deep evolutionary history predating the emergence of the blood circulatory system, as evidenced by its presence in primitive organisms, such as sponges and cnidarians, suggesting an ancient role of locally produced and used C3 within tissues^1^. Research on C3 has traditionally focused on the roles of circulating C3, which is mainly produced by hepatocytes in the liver and is abundantly present in the plasma (100 mg/dL)^4,5^, in host defense mechanisms against infection and various immunoregulatory processes^6,7^. However, recent research has also focused on understanding the functions of locally produced C3 in various biological contexts, such as neurobiology and developmental biology^8,9^, revealing potentially broader roles of C3 than previously appreciated. Despite the growing understanding of local C3 functions, the differential effects of circulating and locally produced C3 on anti-tumor immunity and the efficacy of immune checkpoint blockade (ICB) therapy remain unclear.

Recent studies have demonstrated that C3 is expressed and activated in the cytoplasm of cancer cells, regulating their metabolism and survival^10–12^. Another source of C3 in tumors is cancer-associated fibroblasts (CAFs)^13–15^, which are a major component of the tumor microenvironment (TME) and demonstrate remarkable plasticity in their context-dependent functions and phenotypes^16,17^. Single-cell RNA sequencing (scRNA-seq) analyses have revealed heterogeneous CAF phenotypes, including, but not limited to, α-smooth muscle actin (α-SMA)^+^ and leucine-rich repeat-containing (LRRC) 15^+^ myofibroblastic CAFs (myCAFs) and interleukin (IL)-6^+^ and leukemia inhibitory factor (LIF)^+^ inflammatory CAFs (iCAFs)^18–24^. One of the previously identified CAF subsets is the steady state-like CAFs (sslCAFs)^23,24^, which share characteristics with peptidase inhibitor (PI) 16^+^ tissue-resident “universal” normal fibroblasts that possess intrinsic properties to restrain tumor progression^25,26^ and promote ICB efficacy^27^. A recent study showed that C3 expression is enriched in sslCAFs^23^, implying that C3 may confer on CAFs the ability to promote ICB efficacy through immunoregulatory mechanisms.

We have previously identified Meflin, encoded by the immunoglobulin superfamily containing leucine-rich repeat (ISLR) gene, as a CAF-specific marker expressed in multiple cancer types, including pancreatic^28,29^, colorectal^30^, lung^27^, urothelial^31^, and kidney^31^ cancers. Meflin is a glycosylphosphatidylinositol-anchored membrane protein, and its expression in CAFs ameliorates tissue fibrosis and mechanical stiffening by inhibiting transforming growth factor-β activity via the augmentation of bone morphogenetic protein 7 signaling and reducing collagen cross-linking by inhibiting lysyl oxidase activity^29,30^. Our studies on mouse cancer models and human samples from several cancer types have shown that Meflin plays a cancer-restraining role^27–31^, which is in marked contrast to the standard view that most CAFs promote cancer progression^18–22^. Another feature that distinguishes Meflin from other CAF markers is that Meflin expression enhances ICB efficacy^27,31^, suggesting a potential link between Meflin^+^ CAFs and C3^+^ sslCAFs or universal normal fibroblasts.

In this study, we investigated the role of CAF-derived C3 in modulating the TME and its impact on ICB therapy. Using conditional C3 knockout mouse models and human tumor specimens, we demonstrated that Meflin^+^ CAF-derived local C3, but not systemically circulating C3 produced in the liver, is critical for the anti-tumor efficacy of anti-PD-1 therapy. We showed that C3 expression in CAFs correlated with decreased myeloid cell infiltration into the tumor, leading to a reduction in M2-like tumor-associated macrophages (TAMs) within the TME. Mechanistically, iC3b, a degradation product of CAF-derived C3, inhibited myeloid cell migration by activating complement receptor 3 (CR3), which consists of CD11b (integrin subunit alpha M) and CD18 (integrin subunit beta 2). Finally, we demonstrated that manipulation of CD11b restored anti-PD-1 efficacy in Meflin knockout (KO) mice that were intrinsically resistant to ICB therapy. These results demonstrate the importance of locally produced C3 in the tumor response to ICB, which may be an unexpected benefit of the evolution and development of the complement system.

## RESULTS

### Locally produced C3 determines ICB efficacy

Given that C3 deficiency has been reported to weaken anti-tumor immunity^32–34^ and that approximately 90% of circulating C3 is derived from the liver^4,5^, we first examined the differential roles of systemic and local C3 in anti-tumor immunity. To this end, we used two mouse models, systemic C3 KO mice and hepatocyte-specific C3 KO mice, generated by crossing albumin (*Alb1*)*-Cre* mice with *C3* floxed (*C3*^fl/fl^) mice (Figure S1A–C). C3 expression and its deletion in these mice were confirmed by biochemical and histological analyses, which showed that hepatocyte C3 KO mice lacked C3 expression in the hepatocytes, leading to an 89% reduction in circulating C3 in the plasma (Figure S1D, E). First, we transplanted the MC-38 colon cancer cell line into control, systemic C3 KO, and hepatocyte C3 KO mice and treated them with anti-PD-1 or isotype IgG control antibodies (Figure 1A–C). Using a generalized linear mixed-effects model (GLMM) of tumor growth curves, we evaluated the fixed effects of days post-inoculation (*time*), anti-PD-1 treatment (*P*), systemic C3 KO (*S*), hepatocyte C3 KO (*H*), and their interactions (*time* × *P*, *time* × *S*, *time* × *H*, *time* × *P* × *S*, and *time* × *P* × *H*) on tumor growth rate. The analysis revealed that anti-PD-1 therapy showed significant anti-tumor efficacy (*time* × *P*: estimate -0.547 [95%CI: -0.647; -0.420], P < 0.0001, Figure 1B, C). This effect was markedly attenuated in systemic C3 KO mice completely lacking C3 expression (*time* × *P* × *S*: estimate 0.386 [95%CI: 0.194; 0.578], P = 0.0013, Figure 1B, C) (Figure S1D). In contrast, hepatocyte C3 KO mice maintained anti-PD-1 responsiveness comparable to that of control mice (*time* × *P* × *H*: estimate 0.0322 [95%CI: -0.169; 0.233], P = 0.762, Figure 1B, C). Thus, C3, produced not in the liver but in other sources, plays a major role in the anti-tumor efficacy of anti-PD-1 therapy.

**Figure 1.**
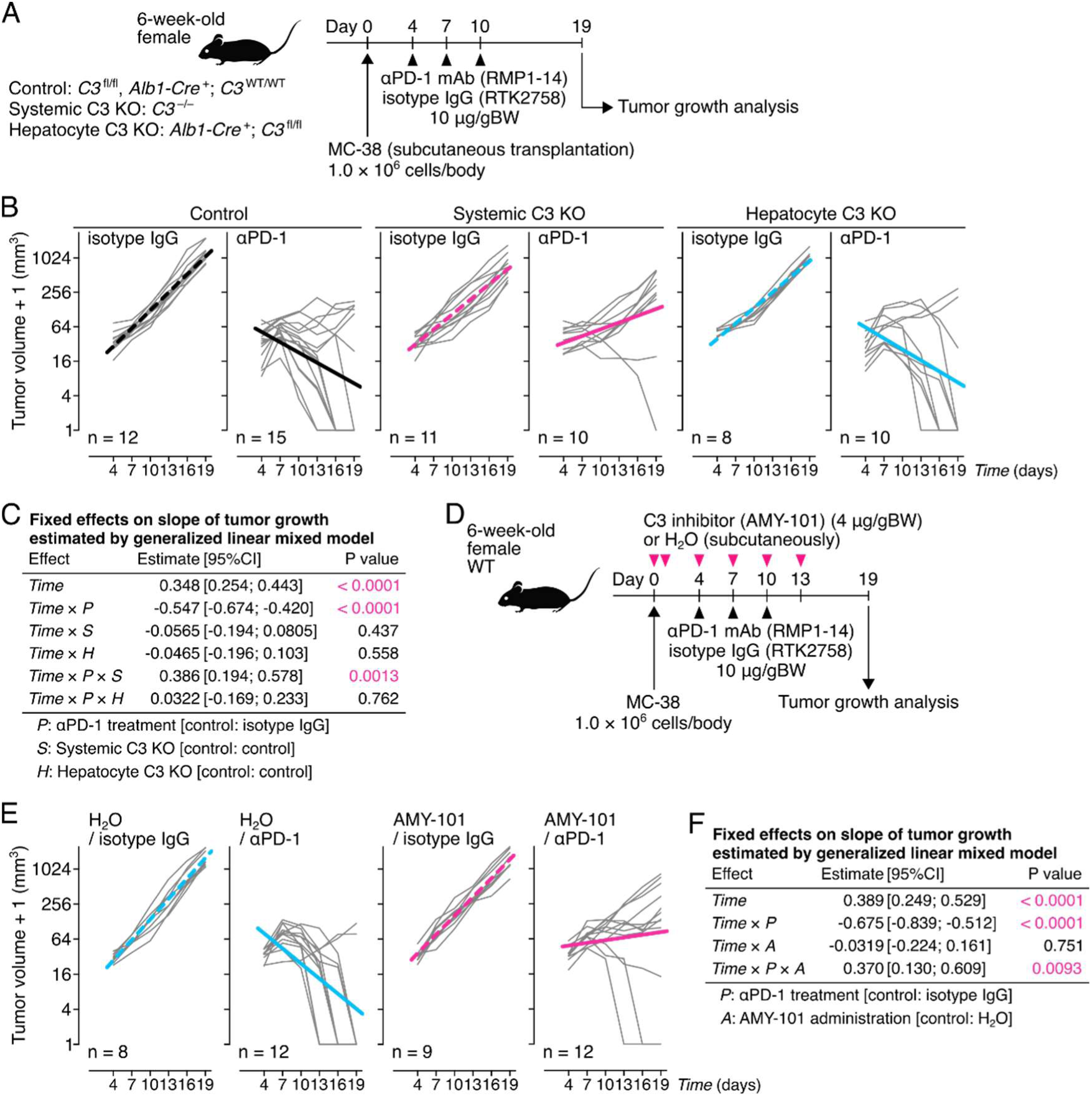
Locally produced, not systemic, complement C3 critically determines the efficacy of ICB therapy. (A) MC-38 mouse colon cancer cells (1.0 × 10^6^ cells) were transplanted subcutaneously into the right dorsal flank of 6-week-old female control (*C3*^fl/fl^ or *Alb1-Cre*^+^; *C3*^WT/WT^), systemic C3 KO (*C3*^-/-^), and hepatocyte C3 KO (*Alb1-Cre*^+^; *C3*^fl/fl^) mice (day 0), followed by intraperitoneal administration of anti-PD-1 (αPD-1) monoclonal antibody (mAb) or isotype IgG at a dose of 10 μg/g body weight (BW). (B) Individual tumor size trajectories (grey lines) with linear approximations (dotted lines: isotype IgG, solid lines: αPD-1, black: control, magenta: systemic C3 KO, cyan: hepatocyte C3 KO) on a log2 scale over 19 days. (C) Fixed effects of days post-inoculation (*time*), αPD-1 (*P*), systemic C3 KO (*S*), hepatocyte C3 KO (*H*), and their interactions on tumor growth rate, estimated using a generalized linear mixed model. Estimates are shown with 95% confidence intervals in square brackets. (D to F), MC-38 cells were transplanted subcutaneously into 6-week-old female wild-type (WT) mice (day 0), followed by intraperitoneal αPD-1 mAb or isotype IgG treatment. The mice also received subcutaneous injections of the selective C3 convertase inhibitor AMY-101 (4 μg/gBW) or water near the tumor site at the indicated times (D), individual tumor size trajectories with linear approximations (dotted lines: isotype IgG, solid lines: αPD-1, cyan: H2O, magenta: AMY-101) (E), and fixed effects of *time*, *P*, AMY-101 (*A*), and their interactions on tumor growth rate estimated using a generalized linear mixed model (F).

Here, we hypothesized that TME-derived local C3 plays a vital role in the efficacy of anti-PD-1 therapy. To investigate the role of local C3, we subcutaneously administered AMY-101^35^, a specific inhibitor of C3 convertase, near the tumor site (Figure 1D–F). While control mice responded significantly to anti-PD-1 therapy, AMY-101 treatment (*A*) markedly reduced this therapeutic effect (*time* × *P* × *A*: estimate 0.370 [95%CI: 0.130; 0.609], P = 0.0093, Figure 1E, F). These complementary genetic and pharmacological approaches demonstrate that local C3 activation in the TME, rather than circulating C3, is necessary for the efficacy of ICB therapy.

### A distinct subset of CAFs expresses C3 in TME

To characterize the CAF subset(s) that express C3 in the TME, we employed publicly available scRNA-seq data from multiple cancer types, including colorectal cancer (CRC), non-small cell lung cancer (NSCLC), and pancreatic ductal adenocarcinoma (PDAC). After standard quality control procedures, we analyzed the stromal cells (95,780 cells) from the three cancer types (Figure 2A). Examination of *C3* and *ISLR* (the gene name for human Meflin) expression revealed that *C3* was primarily expressed in CAFs, particularly in *ISLR*^+^ CAFs (Figure 2B). We validated these findings using single-molecule in situ hybridization (smISH) of tissue samples obtained from patients with colorectal, lung, and pancreatic cancers at our institution. We found a positive correlation between the proportions of *ISLR*^+^ and *C3*^+^ cells among stromal cells, excluding morphologically distinct immune and vascular endothelial cells (Figure S2A–C). Furthermore, single-molecule fluorescent ISH (smFISH) analysis of human lung cancer samples confirmed that C3 expression substantially overlapped with that of *ISLR* (Figure 2C). To further validate C3 expression in *Islr* (the gene name for mouse Meflin)^+^ CAFs in mice, we generated Meflin^+^ CAF-specific C3-deficient (Meflin-lineage C3 KO: hereafter termed mKO) mice by crossing *Islr-Cre* mice with *C3*^fl/fl^ mice (Figure S1C). C3 production was almost undetectable in cultured mouse embryonic fibroblasts (MEFs) isolated from mKO mice, whereas the change in circulating plasma C3 levels was limited to an approximately 9% reduction (Figures 3A and S1D). The number of *Islr*- and *C3*-double-positive CAFs significantly decreased in mT5 and MC-38 tumors that developed in mKO mice compared to those in control mice (Figure S3B, C), supporting the view that Meflin^+^ CAFs produce most of the local C3 in cancer tissue.

**Figure 2.**
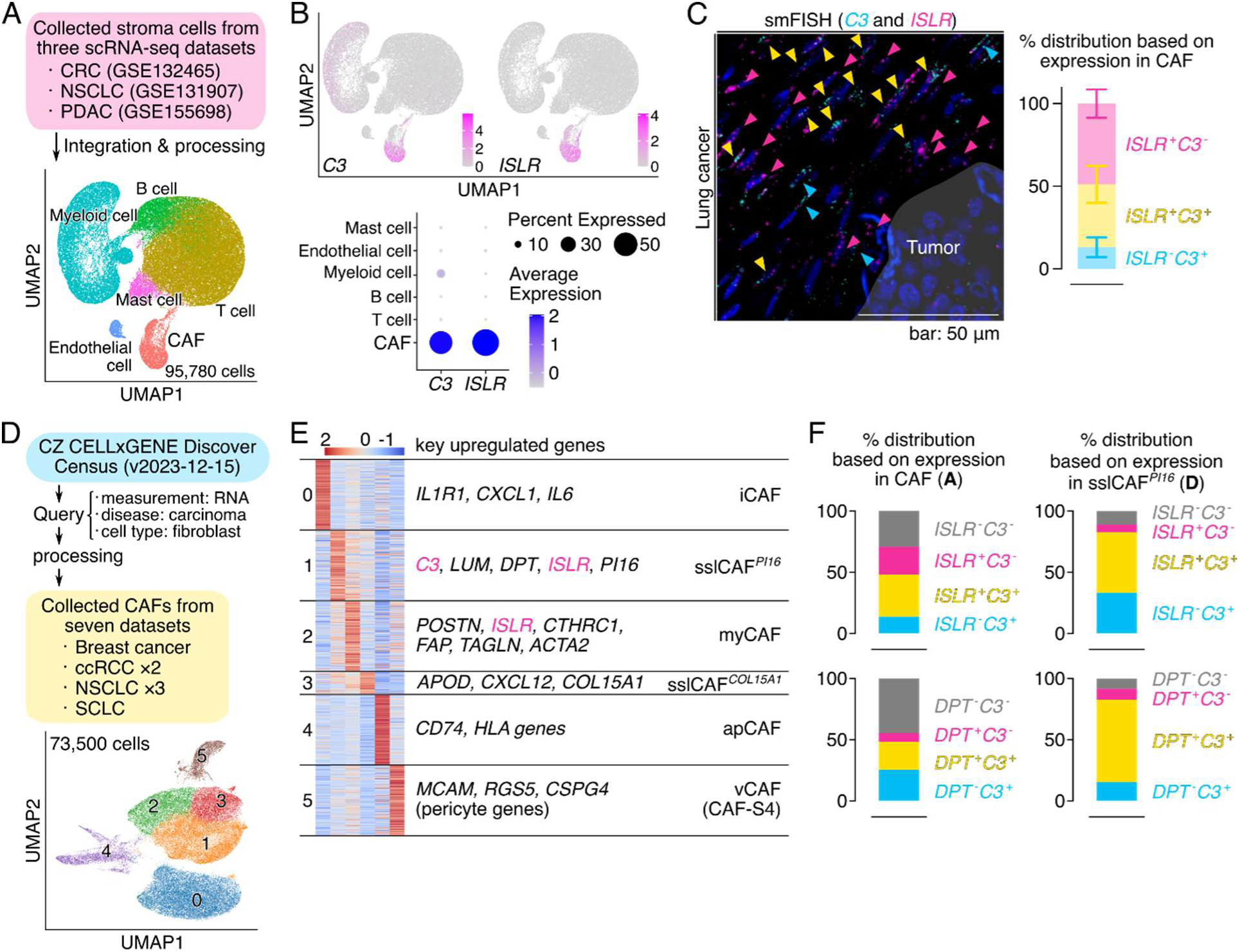
A sslCAF subset expresses C3 in the TME. (A) Uniform Manifold Approximation and Projection (UMAP) visualization of stromal cells from scRNA-seq datasets of colorectal cancer (CRC, GSE132465), non-small cell lung cancer (NSCLC, GSE131907), and pancreatic ductal adenocarcinoma (PDAC, GSE155698). Cells were clustered and colored according to the cell type. (B) Visualization of *C3* and *ISLR* expression levels in stromal cells in the UMAP plot (gray: low/no expression; magenta: high expression) and dot plot. (C) Tissue sections obtained from human NSCLC (n = 6) were stained for *C3* (blue) and *ISLR* (magenta) mRNAs using smFISH. Representative images of *C3* and *ISLR* expression in stromal cells (left) and their mean percentage distributions based on their expression in CAFs (right) (mean ± 95% confidence interval, n = 6). (D) UMAP visualization of CAF subsets generated by Leiden clustering and the CELL×GENE Discover Census database (version 2023-12-15), including breast cancer (n = 1), clear cell renal cell carcinoma (ccRCC, n = 2), NSCLC (n = 3), and small cell lung cancer (SCLC, n = 1). (E) Heatmap of the relative average expression (z-scores) of the most enriched genes (up to 200 genes) for each identified CAF subset: iCAF (*IL1R1* and *IL6*-expressing inflammatory CAFs), sslCAF*^PI16^* (*C3*, *DPT*, *ISLR*, and *PI16*-expressing steady state-like CAFs), myCAF (*POSTN* and *ACTA2*-expressing myofibroblastic CAFs), sslCAF*^COL15A1^* (*APOD* and *COL15A1*-expressing steady state-like CAFs), apCAF (*CD74* and *HLA-DRB1*-expressing antigen-presenting CAFs), and vCAF (*MCAM* and *RGS5*-expressing vascular CAFs or CAF-S4). Red and blue indicate expression levels higher and lower than the mean, respectively, across all clusters. (F) CAFs positive or negative for *ISLR* (upper panels) or *DPT* (upper panels) were classified according to their *C3* expression. The proportions of these populations are shown relative to the total CAF population in panel A and the sslCAF*^PI16^* population (cluster 1) in panel D.

**Figure 3.**
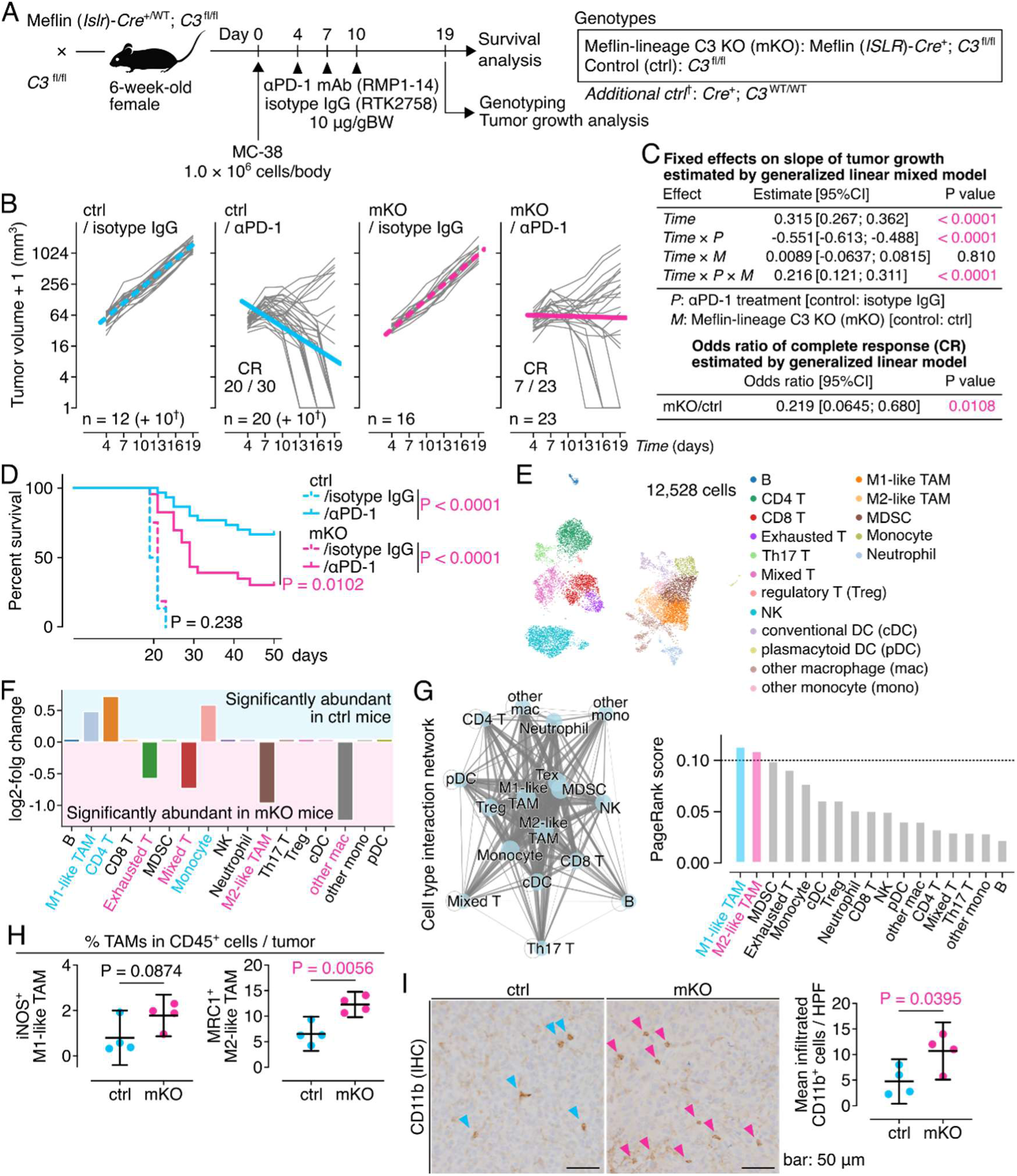
CAF-derived C3 promotes ICB efficacy by limiting the infiltration of immunosuppressive myeloid cells. (A) MC-38 cells were transplanted subcutaneously into 6-week-old female control (*C3*^fl/fl^ and Meflin*-Cre*^+^; *C3*^WT/WT^) or Meflin-lineage C3 KO (mKO; Meflin*-Cre*^+^; *C3*^fl/fl^) mice (day 0), followed by intraperitoneal administration of αPD-1 mAb or isotype IgG. Tumor measurements were performed blind to the genotypes. (B) Individual tumor size trajectories (grey lines) with linear approximations (dotted lines: isotype IgG, solid lines: αPD-1, cyan: control, magenta: mKO) on a log2 scale over 19 days. (C) Fixed effects of days post-inoculation (*time*), αPD-1 (*P*), mKO genotype (*M*), and their interactions on tumor growth rate, estimated using a generalized linear mixed model, and complete response (CR) odds ratio for αPD-1-treated control versus mKO tumors. Estimates and odds ratio are shown with 95% confidence intervals in square brackets. (D) Survival analysis of control and mKO mice treated with the indicated antibodies. Log-rank test P values are shown. (E) UMAP visualization of 12,528 cells from control and mKO tumor samples (n = 2 mice/group). Cells were clustered and colored according to the cell type. (F) scCODA analysis showing significant relative changes (FDR < 0.05) in cell composition between mKO and control tumors. (G) Cell-cell interaction network across cell types (left) and PageRank centrality scores showing M1-like and M2-like TAMs as network hubs (scores > 0.1) (right). (H) Flow cytometric quantification of iNOS^+^ M1-like and MRC1^+^ M2-like TAMs within CD45^+^ cells isolated from control and mKO tumors (gating strategy and representative plots in Figure S6). (I) Total myeloid cells were stained for CD11b by IHC, followed by quantification of the number of CD11b^+^ cells (n = 4 mice, mean of 4 high-power fields [HPFs]/mouse). All analyses were performed on tumors resected on day 9 (E to I).

To further characterize the C3-expressing CAF population, we analyzed 73,500 CAFs from seven datasets spanning four types of cancer: breast cancer (BC), clear cell renal cell carcinoma (ccRCC), NSCLC, and small cell lung cancer (SCLC), followed by clustering into six subsets (Figures 2D, E, and S4A, B). Analysis of the upregulated genes in each cluster revealed that both *C3* and *ISLR* were predominantly expressed in sslCAFs enriched in *PI16* expression (termed the sslCAF*^PI16^* cluster) (Figures 2E, and S4A, B). In the total CAFs and sslCAF*^PI16^* cluster, approximately 50–80% of C3-expressing CAFs co-expressed *ISLR* or *DPT*, which encodes dermatopontin, another marker of sslCAFs or universal normal fibroblasts (Figure 2F)^23,24^. Thus, Meflin^+^ CAFs, which substantially overlap with the previously described sslCAF*^PI16^* subset, represent a source of C3 production in the TME.

### CAF-derived C3 is vital for ICB response

To determine the specific contribution of CAF-derived C3 to the efficacy of immunotherapy, we transplanted MC-38 cells into control and mKO mice. The 6-week-old female mice obtained by crossing mKO and *C3*^fl/fl^ mice were treated with anti-PD-1 or isotype control antibodies according to a predetermined schedule concurrently with tumor measurements and treatments performed in a genotype-blinded manner (Figure 3A–D). Using a GLMM of tumor growth curves, we evaluated the fixed effects of days post-inoculation (*time*), anti-PD-1 treatment (*P*), mKO (*M*), and their interactions (*time* × *P*, *time* × *M*, and *time* × *P* × *M*) on tumor growth rate. Anti-PD-1 therapy was significantly effective in control mice (*time* × *P*: estimate -0.551 [95%CI: -0.613; -0.488], P < 0.0001, Figure 3B, C), but this effect was significantly diminished in mKO mice (*time* × *P* × *M*: estimate 0.216 [95%CI: 0.121; 0.311], P < 0.0001, Figure 3B, C). The complete response rate was also significantly lower in mKO mice (odds ratio: 0.219 [95%CI: 0.0645; 0.680], P = 0.0108, Figure 3B, C), with shortened survival periods following PD-1 blockade therapy (Figure 3D).

These findings demonstrated two contrasting observations: (1) mKO mice exhibited markedly impaired anti-PD-1 responses despite only a 9% reduction in circulating C3, and (2) hepatocyte C3 KO mice maintained anti-PD-1 responses despite lacking 89% of circulating C3. This dissociation between circulating C3 levels and therapeutic efficacy suggests that CAF-derived C3 is a critical determinant of anti-PD-1 response.

### CAF-derived C3 creates a TME conducive to ICB efficacy

Next, we explored how CAF-derived C3 creates an environment conducive to the effectiveness of anti-PD-1 antibodies using several analytical approaches. Bulk RNA sequencing revealed 85 significantly differentially expressed genes in MC-38 tumors that developed in mKO mice compared to those in the control mice (Figure S5A). Gene Ontology over-representation analysis of these genes identified significant enrichment in pathways related to immune cell migration, activation, and adhesion (Figure S5A).

To gain a more detailed understanding of these changes at the cellular level, we performed cellular indexing of transcriptomes and epitopes by sequencing (CITE-seq) of CD45^+^ tumor-infiltrating immune cells to simultaneously profile the expression of cell surface proteins and the whole transcriptome at single-cell resolution. (Figures 3E–G and S5B–F). Initial clustering analysis of 12,528 cells obtained from four individual tumors (Figure S5B, C) successfully identified major immune cell populations, including T cells, B cells, macrophages, dendritic cells, and other immune cells, based on their transcriptional profiles and surface marker expression (Figures 3E and S5D, E). Compositional analysis revealed that the tumors that developed in mKO mice contained significantly more exhausted T cells and M2-like tumor-associated macrophages (TAMs), whereas the tumors that developed in the control mice showed an accumulation of M1-like TAMs and CD4^+^ T cells (Figures 3F and S5F). Network analysis using the PageRank algorithm identified M1-like and M2-like TAMs as crucial cellular hubs for cell-cell interactions between immune cells in the MC-38 model (Figure 3G).

Flow cytometry analysis of cells isolated from MC-38 tumor tissues validated these findings, with M2-like TAMs showing significant accumulation in tumors derived from mKO mice compared to those from control mice (Figures 3H and S6). As TAMs predominantly originate from circulating myeloid cells, including monocytes^36,37^, we further examined the infiltration of a broader range of CD11b^+^ myeloid cells using immunohistochemistry (IHC) of tumor tissue sections (Figure 3I). The results showed a significant increase in myeloid cell accumulation in tumors derived from mKO mice. Thus, CAF-derived C3 limited the accumulation of immunosuppressive M2-like TAMs in the TME by suppressing myeloid cell infiltration, which may enhance the efficacy of anti-PD-1 antibody therapy.

### CAF-derived C3 correlates with improved clinical outcomes

To translate our findings from mouse models to human cancers, we examined the relationship between relative serum complement C3 abundance and treatment outcomes in patients with non-small cell lung cancer (NSCLC) who received ICB therapy at our institute (n = 136, Table S1). The abundances were statistically equivalent between responders and non-responders, with no apparent effect on survival (Figures 4A and S7A).

**Figure 4.**
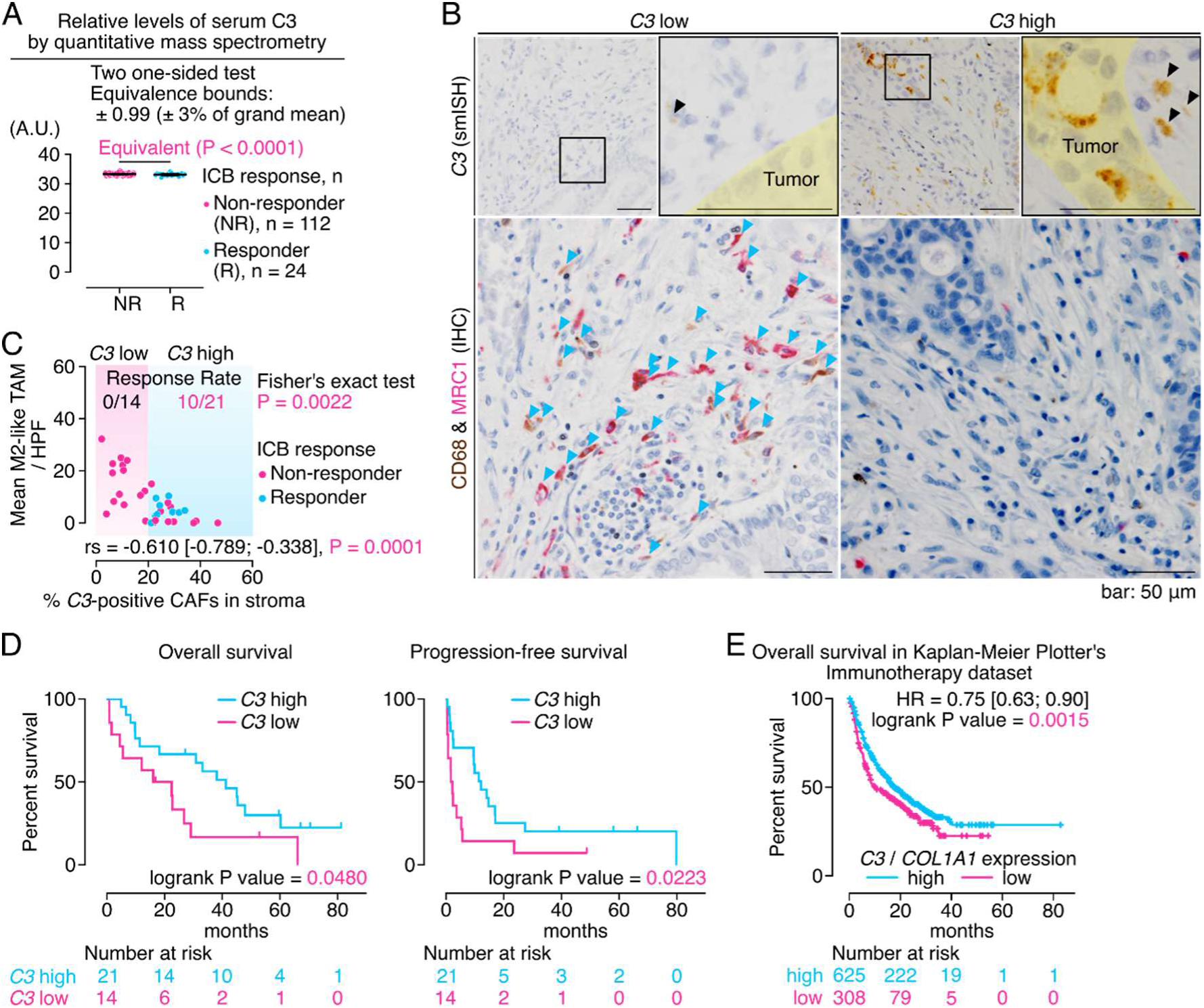
CAF-derived C3, not circulating C3, correlates with reduced M2-like TAM infiltration and improved clinical outcomes in immunotherapy. (A) Relative C3 abundance in the serum of each patient with NSCLC who were responders (R) or non-responders (NR) to ICB therapy was measured using mass spectrometry and showed statistical equivalence between the groups, with equivalence bounds of ± 3%. See Table S1 for patient characteristics. (B) Serial NSCLC tissue sections stained for *C3* by smISH (left) or double-stained for CD68 and MRC1 by IHC (right). Representative images showing cases with high and low proportions of *C3*^+^ CAFs (≥ 20% and < 20%, respectively, referred to as C3 high and C3 low, respectively) in the stroma. Boxed areas are magnified in the lower panels. Yellow: tumor nests. Black and cyan arrowheads indicate *C3*^+^ CAFs and CD68^+^MRC1^+^ M2-like TAMs, respectively. (C) Correlation between the proportion of *C3*^+^ CAFs in the stroma and the number of M2-like TAM infiltrates in tumor tissues from patients with NSCLC, colored by response to ICB therapy (responders: cyan, non-responders: magenta). Each dot represents one patient. Background shading indicates C3 high (≥ 20%, cyan) and low (< 20%, magenta) groups. Response rates for the groups are indicated with Fisher’s exact test P value. Spearman’s rank correlation coefficient (rs) is shown with 95% confidence intervals in parentheses. (D) Kaplan-Meier analysis of overall survival (OS, left) and progression-free survival (PFS, right) comparing C3 high (cyan) and low (magenta) groups. Log-rank test P values are shown. See Table S2 for patient characteristics. (E) Survival analysis of 933 patients with immunotherapy-treated solid tumors from the Immunotherapy dataset in the Kaplan-Meier Plotter. Patients were classified into C3 high (n = 625) and C3 low (n = 308) groups based on stromal *C3* expression normalized to the expression of the fibroblast marker *COL1A1*. Hazard ratio is shown with 95% confidence intervals in square brackets. Number at risk is shown below each survival curve.

We also analyzed stromal C3 expression within the TME and examined its correlation with the number of mannose receptor C-type 1 (MRC1)^+^ M2-like TAMs in NSCLC tissue samples from patients who received ICB therapy (n = 35, Figure S7B, Table S2). The results revealed a significant negative correlation between stromal C3 expression and M2-like TAM infiltration (Spearman’s rank correlation coefficient (rs) = -0.610 [95%CI: -0.789; -0.338], P = 0.0001, Figure 4B, C). Similar negative correlations were observed for colorectal and pancreatic cancers (Figure S8A, B).

To validate these observations, we analyzed scRNA-seq data from colorectal, lung cancer, and pancreatic cancer (Figure 2A) and the Cancer Genome Atlas (TCGA) datasets. Analysis of the scRNA-seq dataset confirmed negative correlations between the mean C3 expression in CAFs and the abundance of both M2-like TAMs and total myeloid cells (Figure S9A, B). A meta-analysis of TCGA datasets, excluding sarcomas (due to the potential of sarcoma tumor cells to express CAF-like genes), brain tumors (where TAMs are predominantly derived from tissue-resident microglia), and hematological malignancies (which lack or contain minimal CAF populations), revealed a negative correlation between stromal *C3* expression (normalized to *COL1A1*) and *MRC1* expression (normalized to *PTPRC*) (Figure S9C).

Using 20% as a threshold for positivity based on previous studies^27,28^, we quantified the number of *C3*^+^ CAFs relative to that of all stromal cells with spindle-shaped nuclei, followed by the categorization of all samples into *C3*^+^ CAF high and low groups (Figure 4B, C). The *C3*^+^ CAF high group had significantly more responders than the low expression group and showed prolonged progression-free and overall survival (Figure 4C, D). These findings were further validated by analyzing a publicly accessible database of ICB-treated patients, with the high stromal *C3* expression group showing a significantly better prognosis (HR= 0.75 [95%CI: 0.63; 0.90], P = 0.0015, Figure 4E). These results in human cancer samples align with our observations in mouse models and suggest the clinical relevance of CAF-derived C3, but not circulating C3, in regulating myeloid cell infiltration and determining ICB therapy outcomes.

### CAF-derived C3 suppresses myeloid cell infiltration through iC3b-CR3 signaling

Based on our observations that CAF-derived C3 inversely correlates with myeloid cell infiltration and considering that M2-like TAMs predominantly arise from infiltrating monocytes^36,37^, we hypothesized that CAF-derived C3 restricts monocyte entry into the TME. To test this hypothesis, we performed transwell migration assays to examine the directional migration of CD11b^+^ cells harvested from the peritoneal cavity in response to serum-free conditioned media collected from cultured control and mKO-derived MEFs. Conditioned media from mKO MEFs significantly enhanced CD11b^+^ cell migration compared to that from control MEF (Figure 5A).

**Figure 5.**
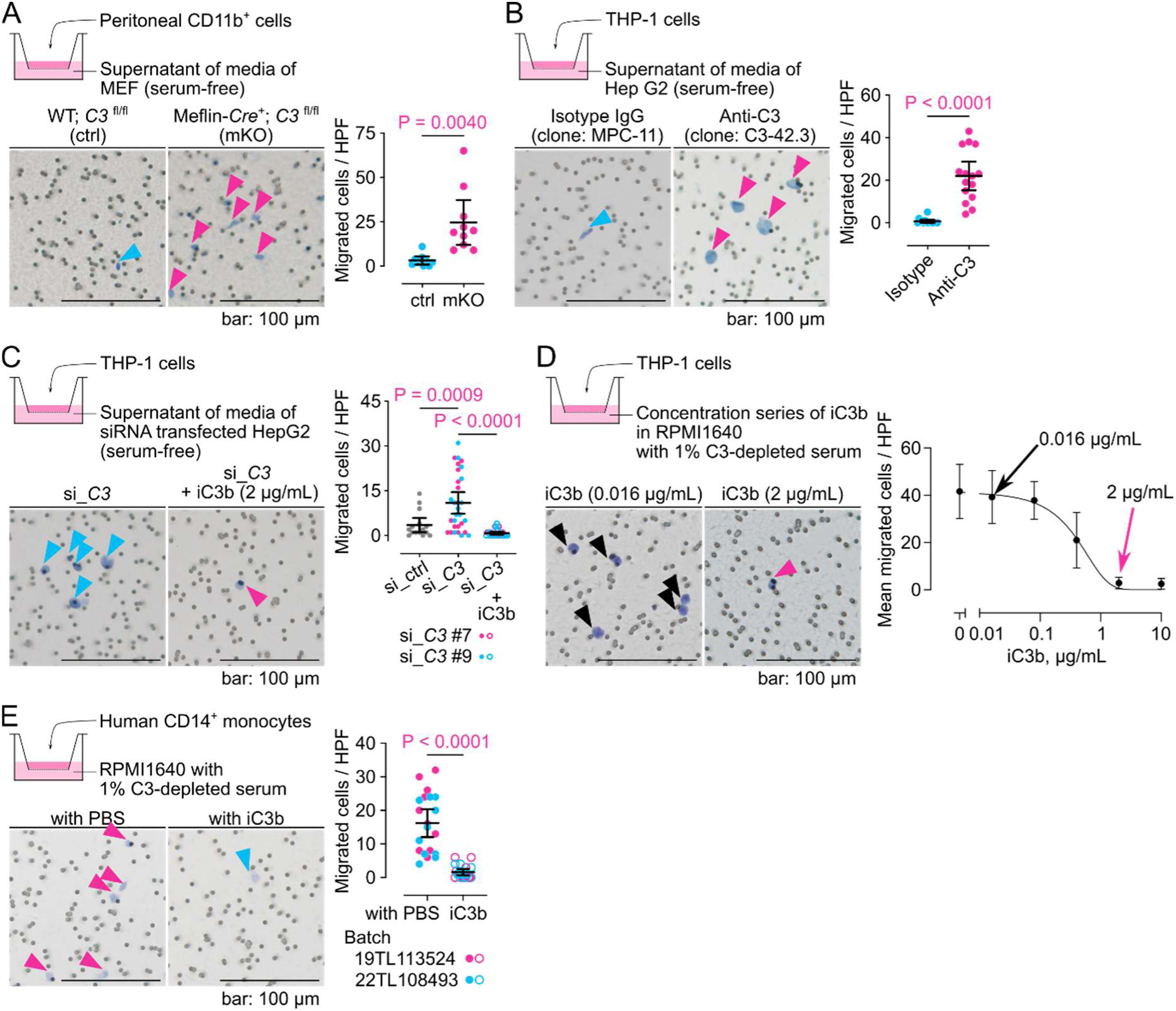
CAF-derived C3 inhibits monocyte migration. (A) Directional migration of mouse peritoneal CD11b^+^ cells in response to serum-free conditioned media from mouse control (*C3*^fl/fl^) and mKO (Meflin*-Cre*^+^; *C3*^fl/fl^) embryonic fibroblasts (MEFs) using transwell assays (n = 2 wells, 5 fields/well). Arrowheads indicate transmigrated cells after 3 hours. (B) Directional migration of THP-1 cells toward HepG2 conditioned medium with anti-C3 or isotype control antibody (1 μM) in the lower chamber, with quantification (n = 3 wells, 5 fields/well). (C) Directional migration of THP-1 cells toward siRNA-transfected HepG2 conditioned medium. The effect of adding purified iC3b (2 μg/mL) to the lower chamber was also examined (n = 3 wells, 5 fields/well). (D) Effects of purified human iC3b (0–10 μg/mL) on THP-1 cell migration in medium containing 1% C3-depleted serum. (E) Directional migration of primary human CD14^+^ monocytes in response to iC3b in medium containing 1% C3-depleted serum (n = 2 donors, 2 wells/donor, 5 fields/well).

We further explored the molecular mechanism underlying C3-mediated inhibition of monocyte migration using the CD11b^+^ human monocytic cell line THP-1 (Figure S10A), and the supernatant from HepG2 hepatoblastoma cells, which are known to produce C3 [38]. Neutralization of C3 with a specific antibody markedly increased THP-1 cell migration compared to isotype control treatment (Figure 5B). Additionally, small interfering RNA (siRNA)-mediated C3 knockdown in HepG2 cells enhanced THP-1 cell migration, an effect reversed by exogenous supplementation with purified human iC3b (Figures 5C and S10B). To further validate the inhibitory effect of iC3b, we performed transwell assays using media containing 1% C3-depleted serum. Purified human iC3b (0.016–10 µg/mL) inhibited THP-1 migration in a dose-dependent manner (Figure 5D). We also confirmed that iC3b inhibited the directional migration of primary human CD14^+^ monocytes, demonstrating the relevance of this regulatory mechanism in human physiology and cancer (Figure 5E). This suppression of monocyte migration was likely mediated by CR3, a heterodimer consisting of CD11b (integrin subunit alpha M) and CD18 (integrin subunit beta 2). Indeed, previous studies have shown that genetic deletion of CD11b results in enhanced myeloid cell infiltration in vivo^39–41^, suggesting that CR3 signaling limits this response.

To establish direct causality, we performed short hairpin RNA (shRNA)-mediated knockdown of CD11b (encoded by the *ITGAM* gene) in THP-1 cells (Figure S10C). CD11b knockdown abolished the inhibitory effect of purified human iC3b on THP-1 cell migration (Figure 6A), confirming that iC3b suppressed monocyte migration via CR3-dependent mechanisms. To delineate the signaling pathway downstream of CR3, we focused on Syk, a key non-receptor tyrosine kinase essential for CR3-dependent responses ^42,43^. Treatment with BAY 61-3606^44^, a selective Syk inhibitor, abolished the anti-migratory effect of iC3b in THP-1 cells (Figure 6B). Notably, neither iC3b nor BAY 61-3606 significantly affected the random migration of THP-1 cells in culture dishes (Figure 6C), suggesting a specific role for the iC3b-CR3 pathway in directional chemotaxis. At the mechanistic level, iC3b treatment significantly reduced THP-1 cell adhesion to the substratum under low-serum conditions, an effect that was reversed by BAY 61-3606 (Figure 6D). These results demonstrate that Syk-dependent signaling mediates the effects of iC3b on both cell adhesion and directional migration. Collectively, our mechanistic studies indicate that CAF-derived C3, through its degradation product iC3b, inhibits monocyte adhesion and migration via the CR3-Syk signaling axis, potentially creating a microenvironment favorable for immune checkpoint blockade therapy.

**Figure 6.**
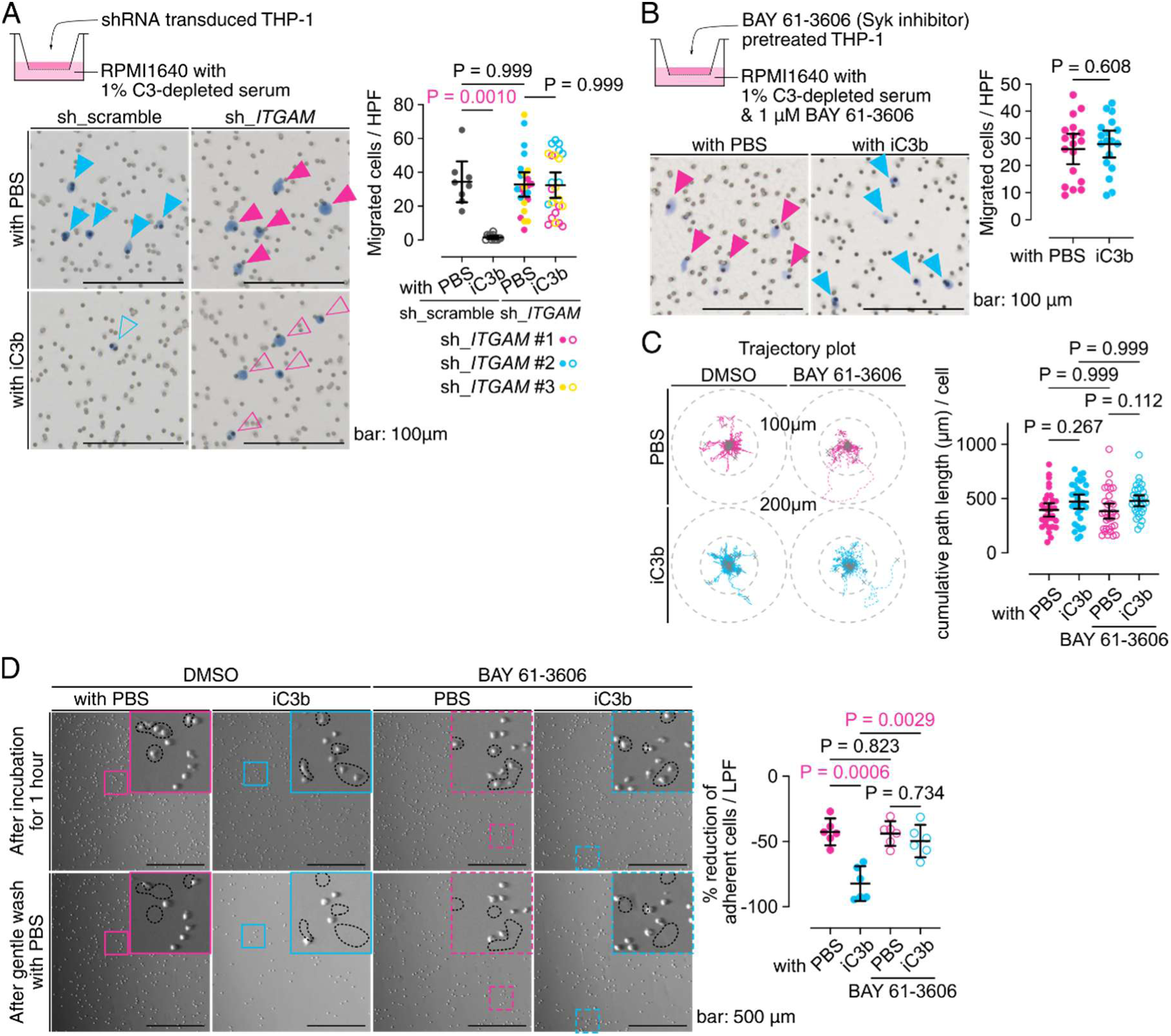
iC3b suppresses monocyte migration via CR3-Syk signaling pathway. (A) Directional migration assay comparing THP-1 cells transduced with control (sh_scramble) or CD11b-targeting (sh_*ITGAM*) shRNAs in response to iC3b in medium containing 1% C3-depleted serum (n = 2 wells, 4 fields/well). Arrowheads indicate transmigrated cells after 3 hours. (B) Effect of selective Syk inhibitor BAY 61-3606 (1 μM) on THP-1 cell migration in response to iC3b in medium containing 1% C3-depleted serum (n = 3 wells, 6 fields/well). (C) Trajectory analysis of THP-1 cells with or without BAY 61-3606 pretreatment in medium containing 1% C3-depleted serum, with or without iC3b (n = 32 cells/group, tracked for 2 hours). (D) Adhesion assay of THP-1 cells cultured on plastic dishes with or without BAY 61-3606 pretreatment, then incubated for 1 hour with or without iC3b in medium containing 1% C3-depleted serum. Images were captured before and after gentle PBS washing to remove non-adherent cells, with quantification of detached cells (n = 2 wells, 3 low-power fields [LPFs]/well). Insets show higher magnification of the boxed areas. Dotted lines indicate regions where cells detached after PBS washing.

### Targeting myeloid cell infiltration restores ICB responsiveness

Our previous studies have demonstrated that Meflin suppresses tissue fibrosis and mechanical stiffening ^28–30^ and confers a cancer-restraining role on CAFs that augments the efficacy of ICB therapy^27,31^. Consistent with this, tumors that developed in Meflin KO mice were intrinsically resistant to ICB therapy and thus can be used as a murine model with stromal defects in sslCAF function^27,31^. We also found that tumors in Meflin KO mice exhibited increased CD11b^+^ myeloid cell infiltration and a significant decrease in the number of C3-expressing CAFs (Figure S11A, B). These data suggest that a deficiency in C3-mediated suppression of CD11b^+^ myeloid cell infiltration contributes to the development of tumors resistant to ICB therapy in Meflin KO mice.

We evaluated the effect of targeting myeloid cell infiltration on the response of tumors that developed in Meflin KO mice to anti-PD-1 therapy (Figure 7A–C). Using a GLMM of tumor growth curves, we evaluated the fixed effects of days post-inoculation (*time*), anti-PD-1 treatment (*P*), CD11b depletion antibody (clone M1/70) (*M*)^45^, CD11b partial agonist ADH-503 (*A*)^46^, and their interactions (*time* × *P*, *time* × *M*, *time* × *A*, *time* × *P* × *M*, and *time* × *P* × *A*) on tumor growth rate. The data showed that anti-PD-1 monotherapy resulted in a significant but weakened response in Meflin KO mice (*time* × *P*: estimate -0.144 [95%CI: -0.222; -0.0664], P = 0.0011, Figure 7B, C). The selective depletion of CD11b^+^ myeloid cells by the administration of CD11b depletion antibody or the suppression of their intratumoral infiltration by the administration of ADH-503 significantly restored the therapeutic sensitivity of MC-38 tumors developed in Meflin KO mice against anti-PD-1 antibody (*time* × *P* × *M*: estimate -0.306 [95%CI: -0.420; -0.193], P < 0.0001, *time* × *P* × *A*: estimate -0.218 [95%CI: -0.329; - 0.106], P = 0.0007, Figure 7B, C). The effect of these combinations on anti-PD-1 therapy efficacy was confirmed by the prolonged survival of Meflin KO mice bearing MC-38 tumors (Figure 7D). Immunophenotyping of tumor tissues confirmed that CD11b-targeting strategies effectively reduced intratumoral CD11b^+^ myeloid cell accumulation (Figure 7E). These findings indicate that the inhibition of CD11b^+^ myeloid cell infiltration restores anti-PD-1 sensitivity in intrinsically or inherently ICB-resistant tumors.

**Figure 7.**
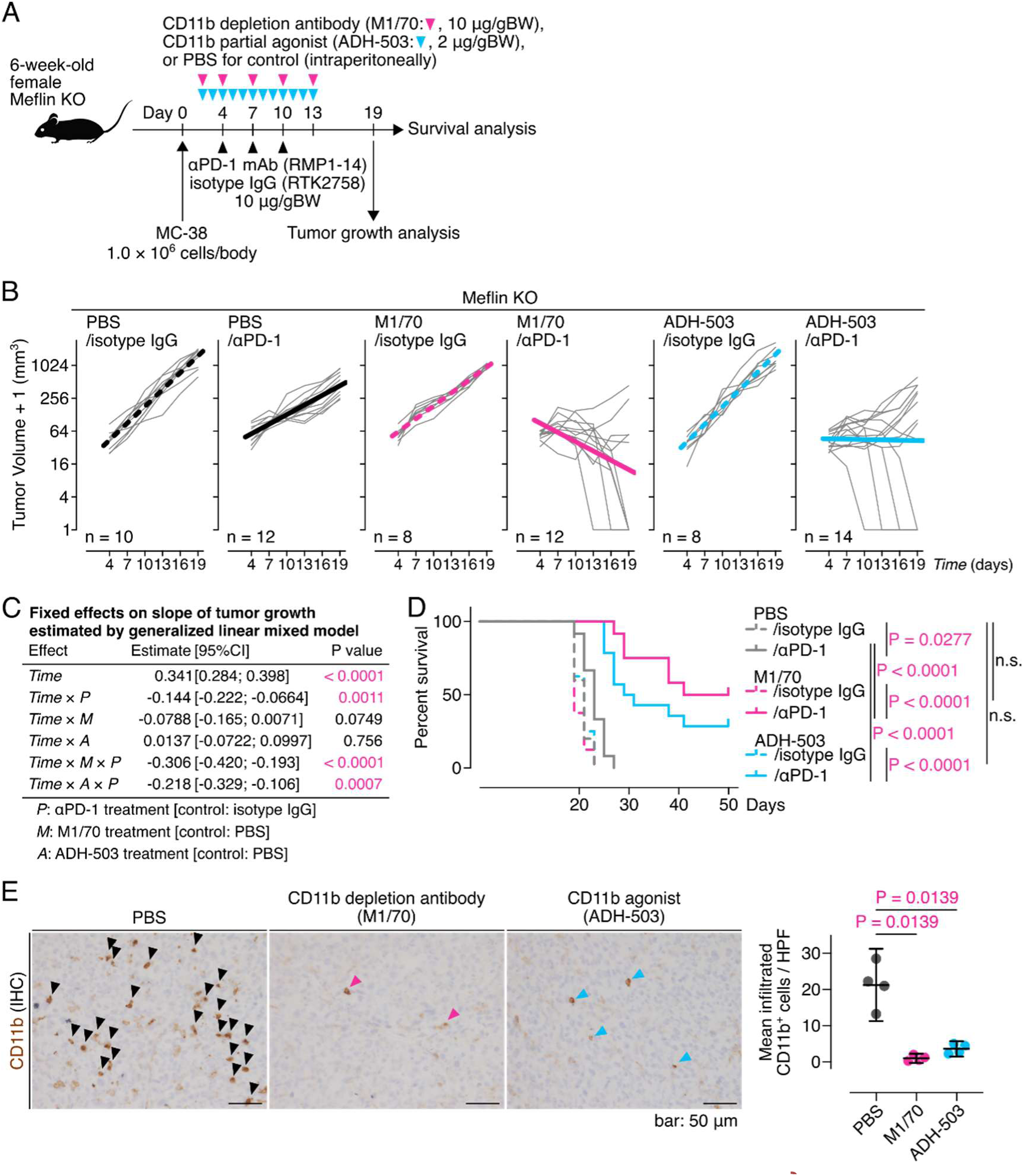
Targeting myeloid cell infiltration restores ICB responsiveness. (A) MC-38 cells (1.0 × 10^6^ cells) were transplanted subcutaneously into 6-week-old female Meflin KO mice (day 0), followed by intraperitoneal αPD-1 mAb or isotype IgG treatment in combinations with CD11b depletion antibody (M1/70: 10 μg/gBW, magenta arrowheads), CD11b partial agonist (ADH-503: 2 μg/gBW, cyan arrowheads), or control (PBS) at the indicated time points. (B) Individual tumor size trajectories (grey lines) with linear approximations (dotted lines: isotype IgG, solid lines: αPD-1, black for PBS, magenta for M1/70, cyan for ADH-503) on a log2 scale over 19 days. (C) Fixed effects of days post-inoculation (*time*), αPD-1 (*P*), M1/70 treatment (*M*), ADH-503 treatment (*A*), and their interactions on tumor growth rate estimated using a generalized linear mixed model. Estimates are shown with 95% confidence intervals in square brackets. (D) Survival analysis of Meflin KO mice treated with the indicated antibodies and agents. Log-rank test P values are shown. (E) Tissue sections from MC-38 tumors developed in Meflin KO mice treated with the indicated agents were stained for CD11b using IHC, followed by quantification of the number of CD11b^+^ cells (n = 4 mice, mean of 4 HPFs/mouse).

## DISCUSSION

Our study revealed two findings that reshaped our understanding of complement function in cancer immunotherapy. First, we demonstrated that local C3, predominantly produced by the sslCAF*^PI16^* subset rather than systemically circulating hepatocyte-derived C3, is a key determinant of the TME and the efficacy of ICB therapy. This finding addresses questions regarding the evolutionary history of C3 and suggests that local and circulating C3s have distinct immunological roles in higher organisms. Second, detailed mechanistic studies demonstrated that CAF-derived C3 creates an immunological niche conducive to ICB therapy by suppressing myeloid cell infiltration through the iC3b–CR3 axis. We also showed that targeting this pathway rendered ICB-resistant tumors sensitive to anti-PD-1 therapy in mice with defective sslCAF function. These findings were reinforced by our clinical analyses, which showed that stromal C3 expression, but not serum C3 levels, was associated with reduced M2-like TAM infiltration and favorable patient responses to ICB and survival across multiple cancer types.

Our findings have several implications for cancer immunotherapy. First, our data showing that the presence of *C3*^+^ CAF subsets from early tumor development is crucial for successful ICB therapy may provide a rationale for developing methods to enhance local C3 production from CAFs or directly or indirectly activate locally produced C3 in the TME rather than systemically modulating complement activity. However, the optimal timing and duration of therapeutic interventions targeting the iC3b–CR3 signaling pathway remain to be determined. This is particularly relevant given that recent clinical studies have shown the failure of agents targeting myeloid cells, such as ADH-503 (NCT04060342), in established and advanced tumors^47^. Second, the significant correlation between stromal C3 expression and clinical outcomes suggests its potential as a predictive biomarker for the clinical benefits of ICB therapy. Therefore, the development of standardized methods for assessing stromal C3 expression and establishing validated cutoff values across different cancer types are required for successful clinical translation.

The molecular mechanisms controlling C3 expression in specific CAF subsets require further investigation. Although we identified the sslCAF*^PI16^* subset as the primary source of C3, the factors and mechanisms that maintain the phenotype of sslCAFs and their plasticity to switch between different types of CAFs are yet to be investigated. The balance between transforming growth factor-β (TGF-β) and retinoic acid (RA) signaling may be crucial in regulating the sslCAF phenotype, as our previous studies showed that Meflin expression was significantly attenuated by TGF-β whereas augmented by RA^29,30^. A better understanding of the local or systemic cues that regulate C3 expression in the sslCAF*^PI16^* subset may help to identify new therapeutic targets. The mechanism by which tumor cells counteract CAF-derived C3 to suppress anti-tumor immunity also needs to be further studied. A recent study showed that tumor cells upregulate the C3 convertase inhibitors CD55 and CD59, which is one of the mechanisms by which tumor cells shape the ICB-resistant TME^48^.

Local C3 production by fibroblasts within tissues has also been implicated in other inflammatory conditions. For example, tissue-resident fibroblasts express C3 and undergo priming in response to inflammation, which is essential for either sustained inflammation or tissue repair, depending on the context^49,50^. This is the first study to demonstrate the role of locally produced C3 in disease conditions. Further studies on the role of local C3 in other biological contexts will improve our understanding of the benefits of the innate immune system, which is conserved from very primitive organisms to humans.

## RESOURCE AVAILABILITY

### Lead contact

Further information and requests for resources and reagents should be directed to and will be fulfilled by the lead contact, Atsushi Enomoto.

### Materials availability

The mouse line generated in this study will be deposited to RIKEN BRC and will be available for non-commercial research purposes with a completed materials transfer agreement.

### Data and code availability

- Bulk RNA-seq and CITE-seq data have been deposited in the Genomic Expression Archive (GEA) under accession numbers E-GEAD-1048 and E-GEAD-1049 and are publicly available as of the date of publication.
- This paper does not report the original code.
- Any additional information required to reanalyze the data reported in this paper is available from the lead contact upon request.

## ACKNOWLEDGMENTS

This study was funded by the Japan Agency for Medical Research and Development through grants JP24gm1210009, JP24ama221333 (to A.E.); the Ministry of Education, Culture, Sports, Science, and Technology of Japan through grants 23K14591, 25K18851 (to Y.M.), 22H03350, 22H04923 (to M.N.), 20H03528 (to Y.A.), and 22H02848, 22K18390 (to A.E.); Aichi Cancer Research Foundation (to Y.M.); the Naito Foundation (to A.E.); Princess Takamatsu Cancer Research Fund (to A.E.); DAIKO Foundation (to A.E.); and the Toyoaki Foundation (to A.E.). We acknowledge the European Conditional Mouse Mutagenesis Program (EUCOMM) Consortium as the source of the targeted embryonic stem (ES) cell clones (HEPD0812_4_D01, E04, and H02) used to generate the conditional C3 knockout mouse model in this study. We thank Dr. Derek LeRoith (Icahn School of Medicine at Mount Sinai), Dr. Takayoshi Suganami, and Dr. Miyako Tanaka (Research Institute of Environmental Medicine, Nagoya University) for providing the hepatocyte-specific Alb1-Cre mouse line; David Tuveson (Cold Spring Harbor Laboratory) and Chang-il Hwang (UC Davis College of Biological Sciences) for providing the mouse PDAC cells mT5; Fumiya Ito (Nagoya University Graduate School of Medicine) for providing the CD11b^+^ monocytic cells THP-1; Kozo Uchiyama, Eri Yorifuji (Nagoya University), and Yukie Atsumi (AGC Inc.) for technical assistance; Rika Fritsch (Medical University of Vienna) for experimental assistance; Yasuko Sobue (Nagoya University) for maintaining our animal colonies. We would like to thank Editage (www.editage.jp) for English language editing.

## AUTHOR CONTRIBUTIONS

Conceptualization, Y.M.; methodology, Y.M., Yu.S., D.S., Yo.S., K.K., N.A., N.N., and K.T.; Investigation, Y.M., Yu.S., C.F., N.N., K.T., and M.N.; Writing – original draft, Y.M. and A.E.; Writing – review & editing, Y.M., Yu.S., N.N., K.T., M.N., N.E., S.M., and A.E.; Funding acquisition, Y.M., M.N., Y.A., and A.E.; Resources, Yu.S., T.H., T.F.C-Y., T.K., S.I., and H.A.; Supervision, H.K., M.T., Y.A., and A.E.

## DECLARATION OF GENERATIVE AI AND AI-ASSISTED TECHNOLOGIES

During the preparation of this work, the authors used Claude in order to improve readability and language. After using this tool or service, the authors reviewed and edited the content as needed and take full responsibility for the content of the publication.

## SUPPLEMENTAL INFORMATION

**Figure S1.**
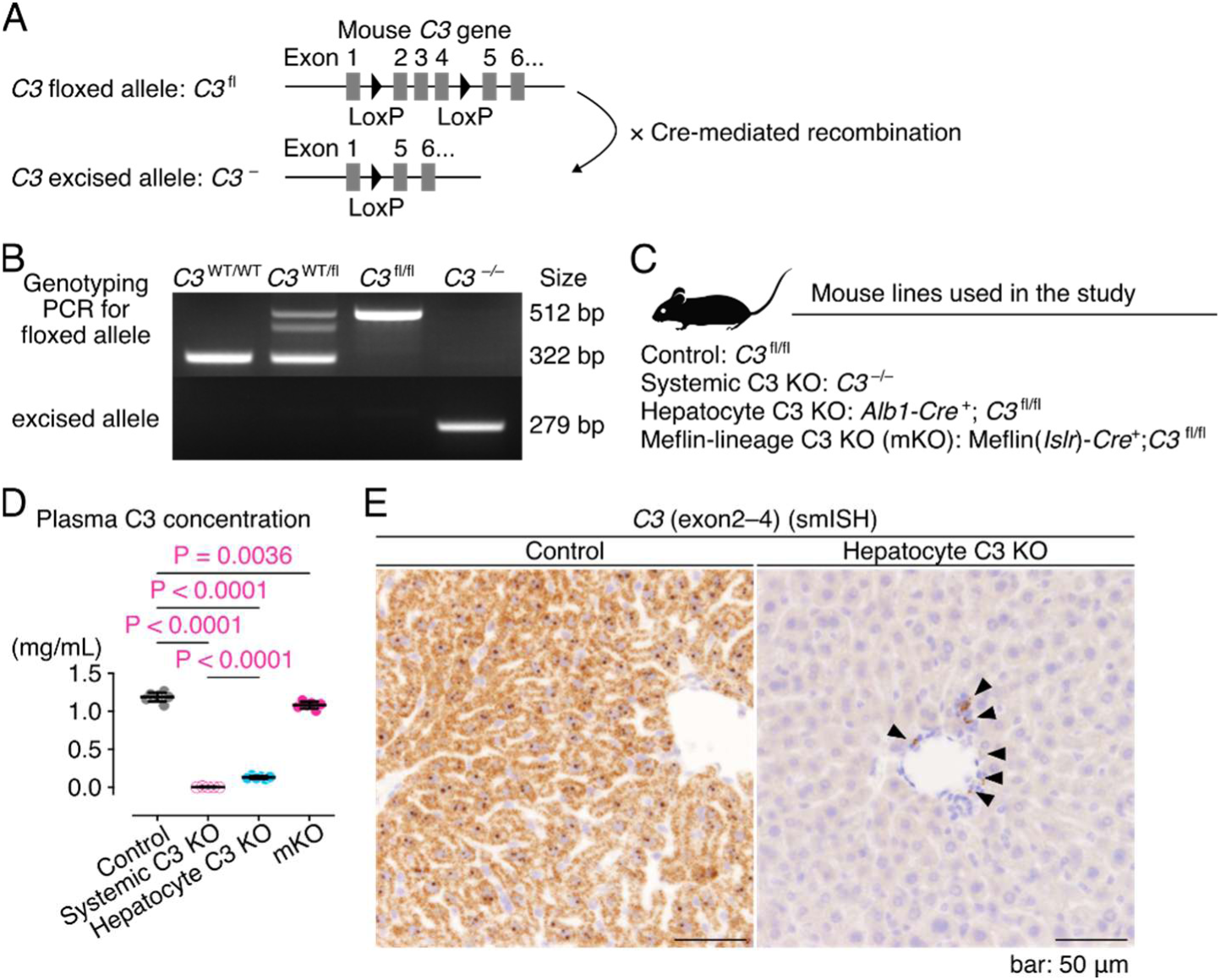
Generation and validation of C3 conditional knockout mouse models and analysis of C3 levels in the plasma and liver. (A) Schematic diagram of the C3 floxed allele (upper panel) and its excision after Cre-mediated recombination (lower panel). In the C3 floxed allele, exons 2–4 were flanked by LoxP sites. (B) Representative PCR genotyping data showing alleles of wild-type (322 bp), heterozygous floxed (*C3*^WT/fl^, 322/512 bp), homozygous floxed (*C3*^fl/fl^, 512 bp), and systemic knockout (*C3*^-/-^, 279 bp) mice. (C) Mouse lines used in this study: control (*C3*^fl/fl^), systemic C3 knockout (KO) (*C3*^-/-^), hepatocyte C3 KO (*Alb1-Cre*^+^; *C3*^fl/fl^), and Meflin-lineage C3 KO (mKO: Meflin (*Islr*)*-Cre*^+^; *C3*^fl/fl^). (D) The plasma C3 concentrations measured using ELISA (n = 5 per group). (E) *C3* mRNA expression in liver sections of control (left) and hepatocyte C3 KO mice (right) was examined through smISH using a probe targeting exons 2–4. Strong positive signals (brown) were observed throughout the livers of control mice, whereas *C3* expression was limited to interstitial cells in the livers of hepatocyte C3 KO mice (arrowheads).

**Figure S2.**
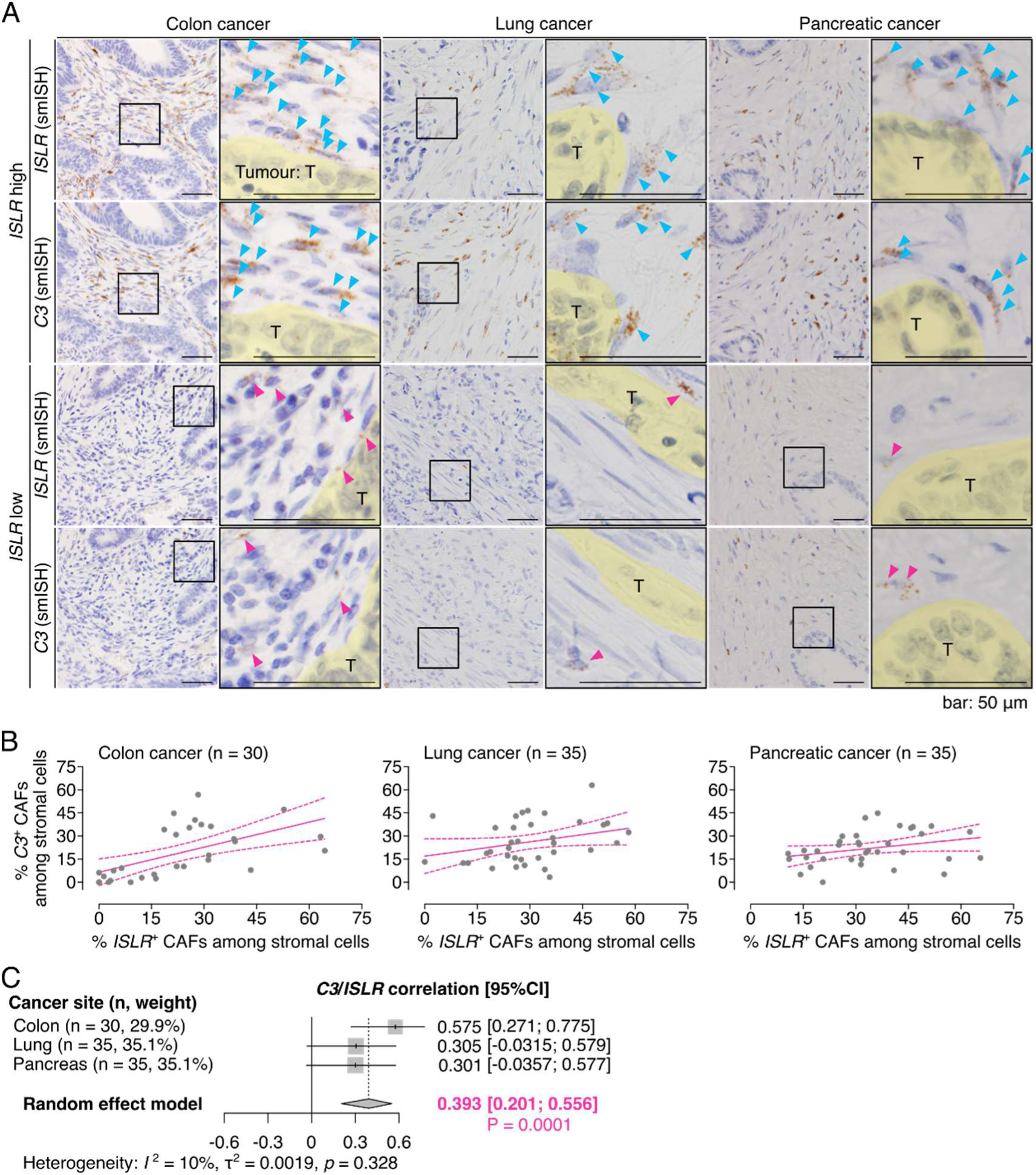
Correlation between *ISLR* and *C3* expression in human cancers. (A) Representative smISH images of serial sections showing spatial distribution patterns of *ISLR* and *C3* expression in human colorectal (left), lung (middle), and pancreatic (right) cancer tissues. Representative data from patients with high (upper panels) and low (lower panels) proportions of *ISLR*^+^ CAFs in the stroma. The boxed areas are magnified in the adjacent panels. Arrowheads indicate smISH-positive signals. T, tumor. (B) Scatter plots illustrating the quantitative correlation between *ISLR* and *C3* expression levels determined by image analysis of smISH data from panel A. Each dot represents the proportion of *ISLR*^+^ and *C3*^+^ stromal cells present in a high-power field (x400) with fitted regression lines (solid lines) and their 95% confidence intervals (dotted lines). (C) Forest plot displaying the results of the meta-analysis of *C3*-*ISLR* expression correlation across the three cancer types. Each data point represents Pearson’s correlation coefficient with a 95% confidence interval.

**Figure S3.**
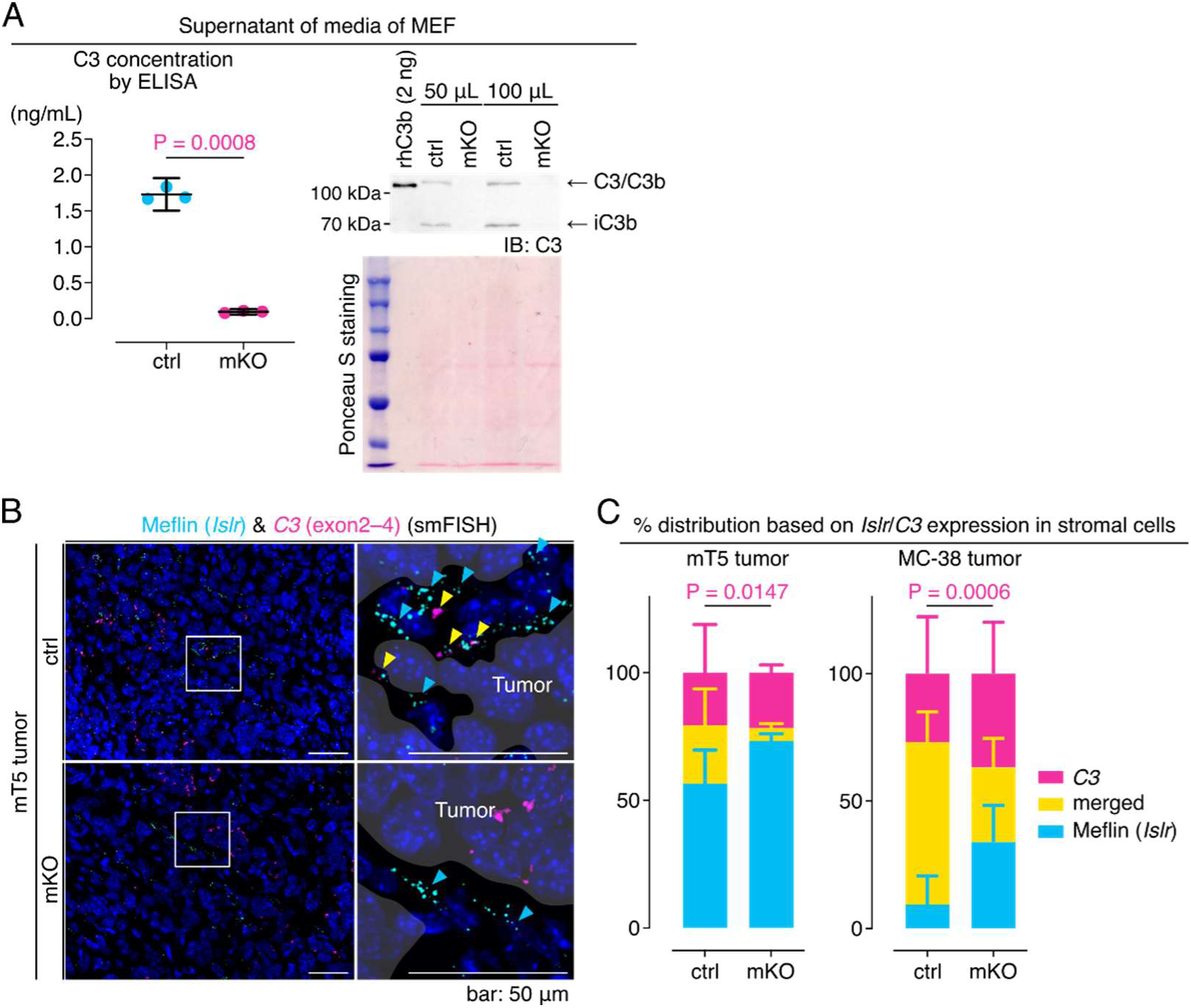
Meflin^+^ fibroblasts are a source of C3. (A) Analysis of mouse embryonic fibroblast (MEF) media supernatants derived from control (ctrl) and Meflin-lineage C3 KO (mKO) mice. Detection of C3 in the media supernatants using ELISA (left) and immunoblotting (IB) (right). Recombinant human C3b protein (2 ng) served as positive control. The α-chain of C3/C3b (115 kDa) and α-chain fragment of iC3b (68 kDa) were detected by IB using an anti-C3 antibody (upper right). Ponceau S staining indicated an equal loading of total proteins (lower right). (B) Tissue sections from mT5 tumors developed in control (upper panels) and mKO (lower panels) mice were stained for Islr (cyan) and C3 (exons 2–4, magenta) using smFISH. The boxed areas are magnified in the adjacent panels. Yellow arrowheads denote Islr and C3 double-positive cells, which were not observed in tumors that developed in mKO mice. (C) Percentage distribution of stromal cells based on *Islr* and *C3* expression in mT5 (left) and MC-38 (right) tumors developed in control and mKO mice.

**Figure S4.**
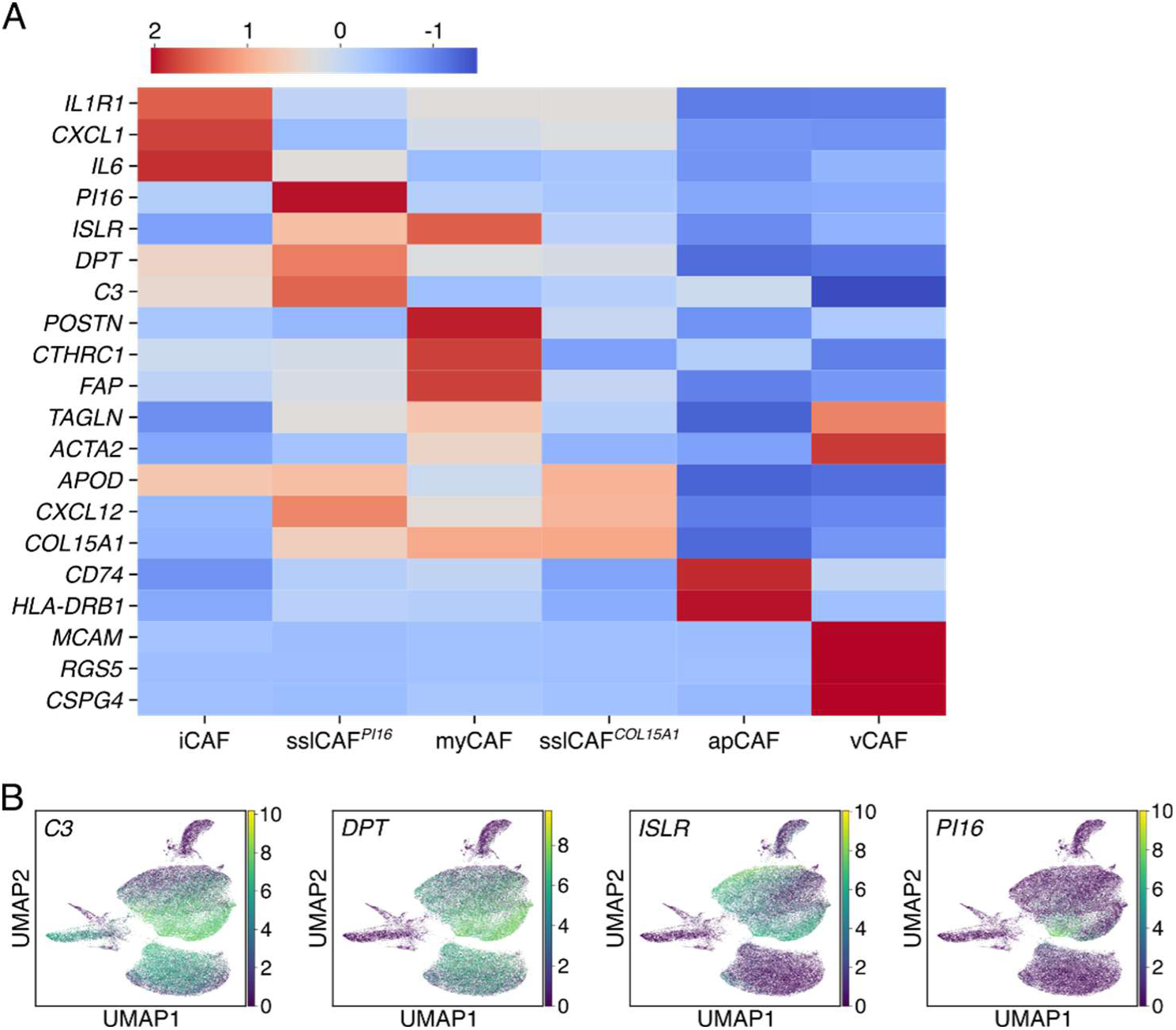
Single-cell RNA sequencing analysis of cancer-associated fibroblasts. (A) Heatmap of the relative average expression (z-scores) of the key genes listed in Figure 2F enriched in iCAF, sslCAF*^PI16^*, myCAF, sslCAF*^COL15A1^*, apCAF, and vCAF. Red and blue indicate higher and lower expression levels than the mean across all clusters, respectively. (B) The expression of *C3*, *DPT*, *ISLR*, and *PI16* in CAFs was visualized using UMAP plots (purple: low/no expression; yellow: high expression).

**Figure S5.**
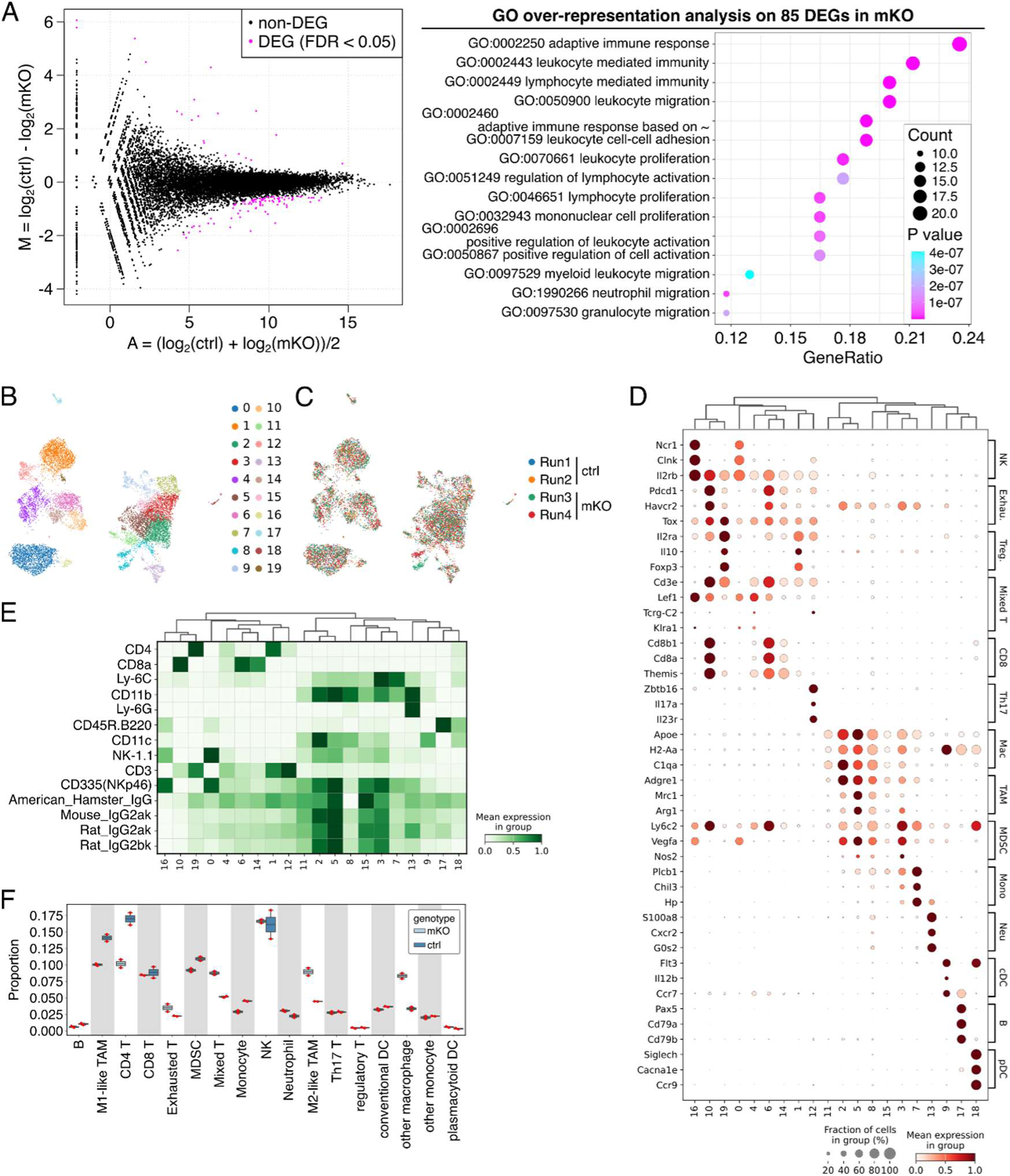
Bulk RNA-seq and CITE-seq analyses of tumors developed in control and mKO mice. (A) Bulk RNA-seq analysis of control and mKO MC-38 tumor samples was visualized using an MA plot to show differentially expressed genes (DEGs, FDR <0.05) between groups (left). Gene Ontology over-representation analysis of the 85 upregulated DEGs identified in mKO tumors (right). (B, C) UMAP visualizations of distinct cell clusters identified in the CITE-seq data using the Leiden clustering algorithm (B), cells from four individual tumor samples (Runs 1–4) with genotype (control and mKO) to assess batch effects (C). (D) Dot plot depicting the expression of the three marker genes for each cell type across the Leiden clusters. (E) Heatmap showing cell surface antigen expression levels measured by CITE-seq across Leiden clusters. (F) Bar plot comparing the proportions of the indicated cell types identified in MC-38 tumors between the control and mKO mice. Bars represent the mean values of two experimental samples. Red dots show the actual values for each sample.

**Figure S6.**
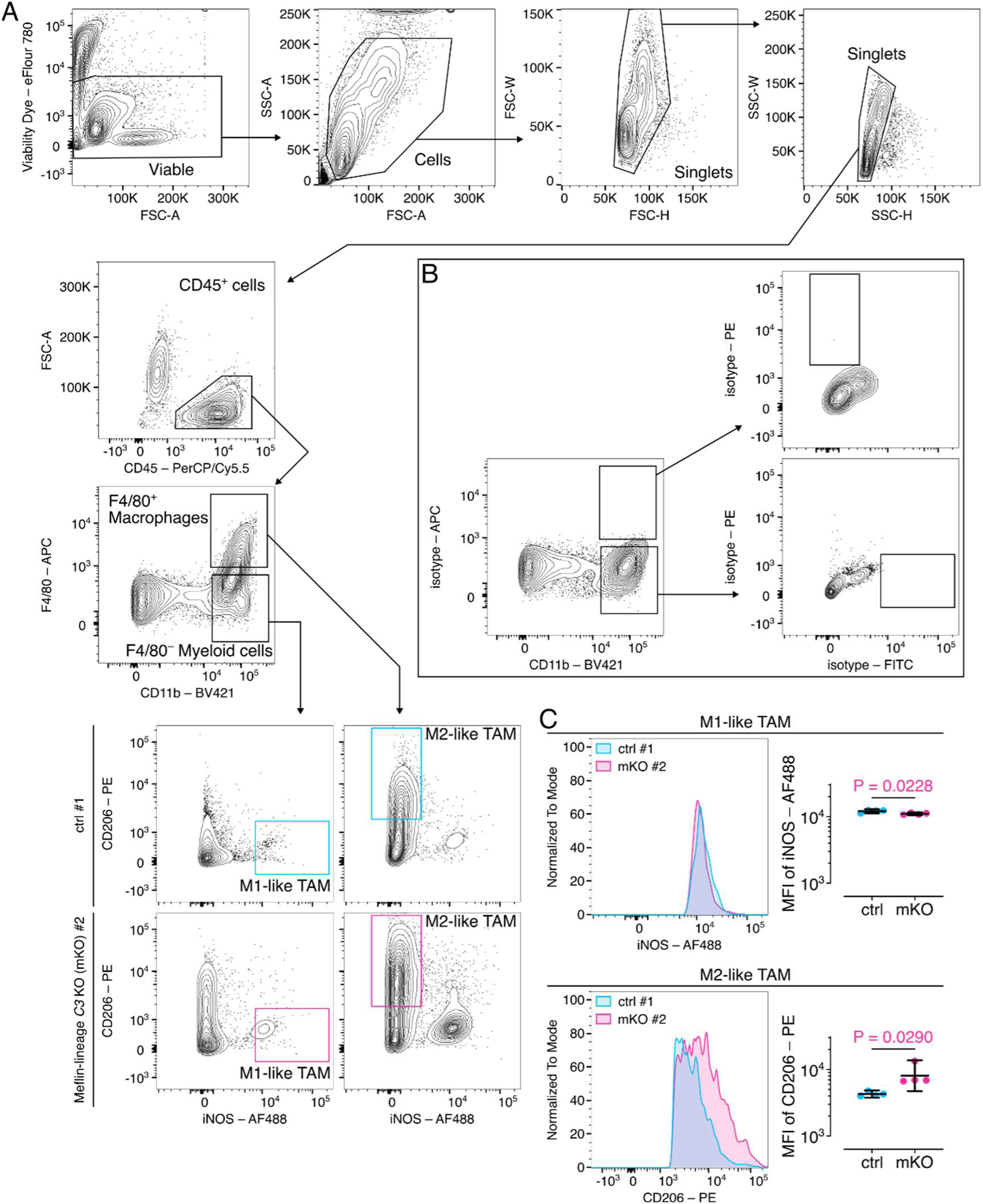
Flow cytometry analysis of tumor-associated macrophages. (A) Flow cytometry gating strategy for identifying TAMs within the total cells of MC-38 tumors developed in control (cyan) and mKO (magenta) mice. (B) Representative flow cytometry plots showing negative control experiments using isotype control antibodies against F4/80–APC, iNOS–AF488, and CD206–PE. (C) Representative histograms showing the mean fluorescence intensity (MFI) of iNOS in gated CD11b^+^F4/80^-^ cells and CD206 in gated CD11b^+^F4/80^+^ cells isolated from MC-38 tumors developed in control (cyan) and mKO (magenta) mice and quantified from four independent experiments.

**Figure S7.**
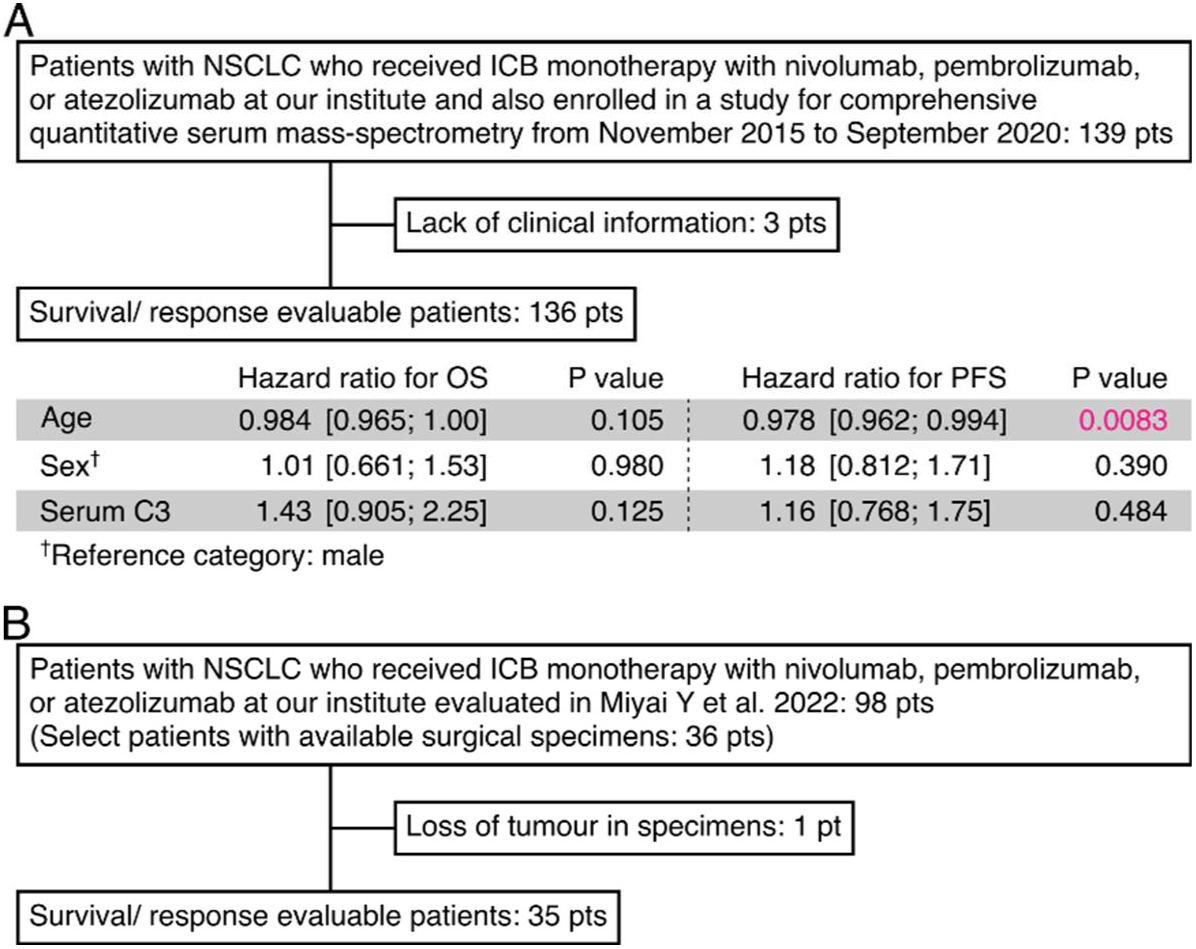
Selection of patient cohorts for the analysis of serum C3 concentration and tissue C3 expression. (A) Flow diagram and multivariate analysis of 136 patients with NSCLC who received ICB monotherapy (nivolumab, pembrolizumab, or atezolizumab) and were enrolled in the comprehensive quantitative serum mass spectrometry analysis from November 2015 to September 2020. The associated table shows the multivariate analyses of overall survival (OS) and progression-free survival (PFS), including hazard ratios (HRs) with 95% confidence intervals for age, sex (reference: male), and serum C3 levels. (B) Flow diagram shows the selection of 98 patients with NSCLC who received ICB monotherapy, with 35 patients having available surgical specimens.

**Figure S8.**
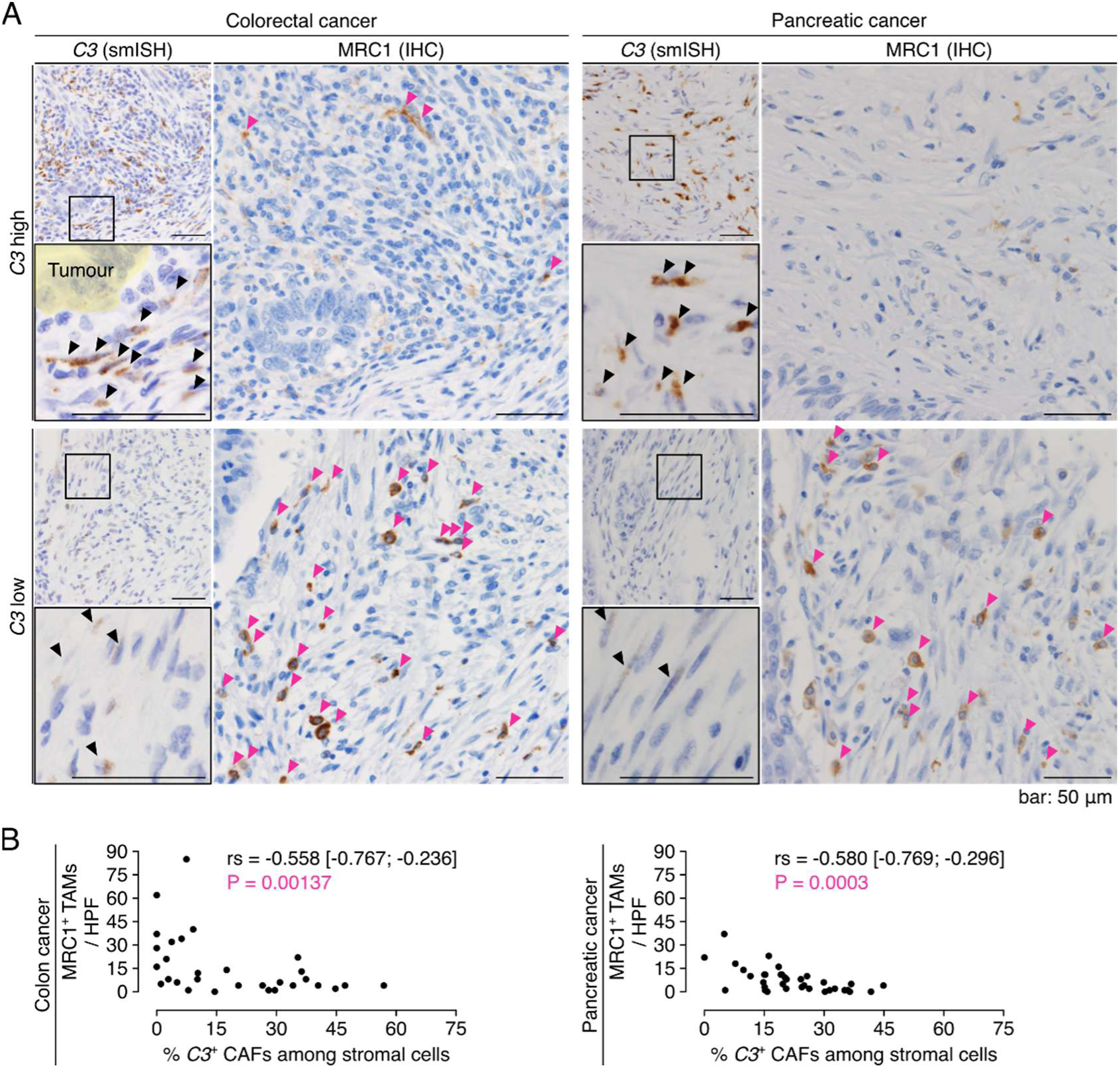
Correlation between C3 expression in CAFs and MRC1+ TAM infiltration in colon and pancreatic cancer. (A) Serial tissue sections obtained from patients with colon (left) and pancreatic (right) cancers were stained for *C3* by smISH or MRC1 by IHC. Representative images of cases with high (C3 high, upper panels) and low (C3 low, lower panels) proportions of *C3*^+^ CAFs in the stroma. Boxed areas are magnified in the lower panels. Yellow shaded areas represent tumor cell nests. The black and red arrowheads indicate *C3*^+^ CAFs and MRC1^+^ M2-like TAMs, respectively. (B) Scatter plots showing the correlation analysis between the *C3*^+^ CAF proportion in the stroma and MRC1^+^ M2-like TAM infiltration in tumor tissues obtained from patients with colon (left) and pancreatic (right) cancers. Each dot represents data from a high-power field (HPF). Statistical analysis was performed using Spearman’s rank correlation coefficient (rs).

**Figure S9.**
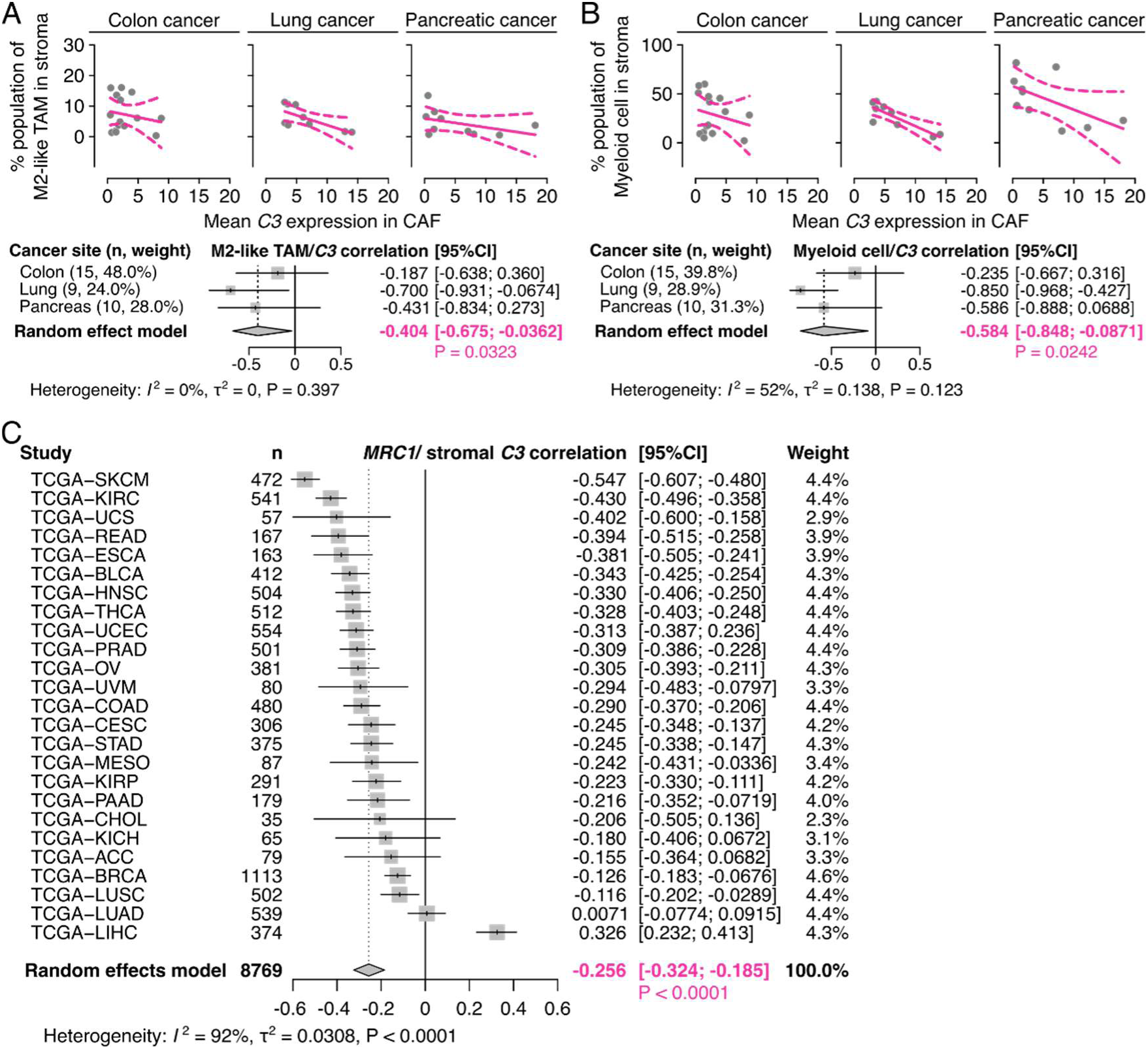
Correlation between stromal C3 expression and MRC1+ TAM infiltration in human solid cancers. (A, B) Analysis of publicly available scRNA-seq datasets from colorectal (GSE132465), lung (GSE131907), and pancreatic (GSE155698) cancers (Figure 2A). Correlations between mean *C3* expression in CAFs and tumor-infiltrating M2-like TAM (A) or total myeloid cell (B) proportions are shown as scatter plots. Each dot represents a patient. Fitted regression lines with 95% confidence intervals are shown for each cancer type. Forest plots of the meta-analysis results showing correlations across cancer types are shown as scatter plots. Pearson’s correlation coefficients with 95% confidence intervals are shown. (C) Meta-analysis results of the correlations between *C3* expression (normalized to *COL1A1*) and *MRC1* expression (normalized to *PTPRC*) in various human solid cancers using the TCGA database, excluding sarcomas, brain tumors, and hematological malignancies.

**Figure S10.**
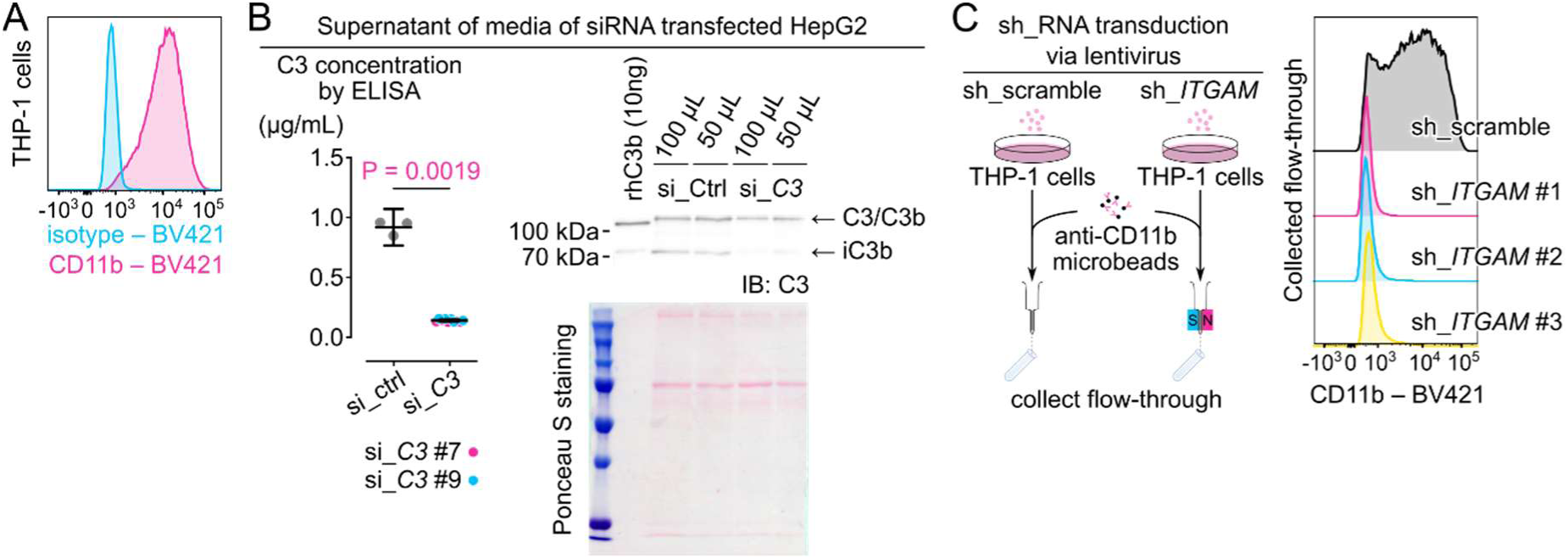
Analysis of CD11b and C3 in THP-1 and HepG2 cell. (A) Flow cytometric analysis of THP-1 cells for CD11b expression using BV421-conjugated antibodies. (B) Detection of C3 in siRNA-transfected HepG2 cell supernatants by ELISA (left) and immunoblotting (right). Recombinant human C3b protein (10 ng) served as a positive control. The α-chain of C3/C3b (115 kDa) and the α-chain fragment of iC3b (68 kDa) were detected by IB using an anti-C3 antibody (upper right). Ponceau S staining indicated an equal loading of total proteins (lower right). (C) Characterization of shRNA-mediated CD11b (*ITGAM*) knockdown in THP-1 cells. Cells were transduced with either control (sh_scramble) or CD11b-targeting (sh_*ITGAM*) shRNAs, with sh_ITGAM-transduced cells further enriched via MACS using anti-CD11b microbeads. CD11b expression in both control and knockdown cells was assessed by flow cytometry.

**Figure S11.**
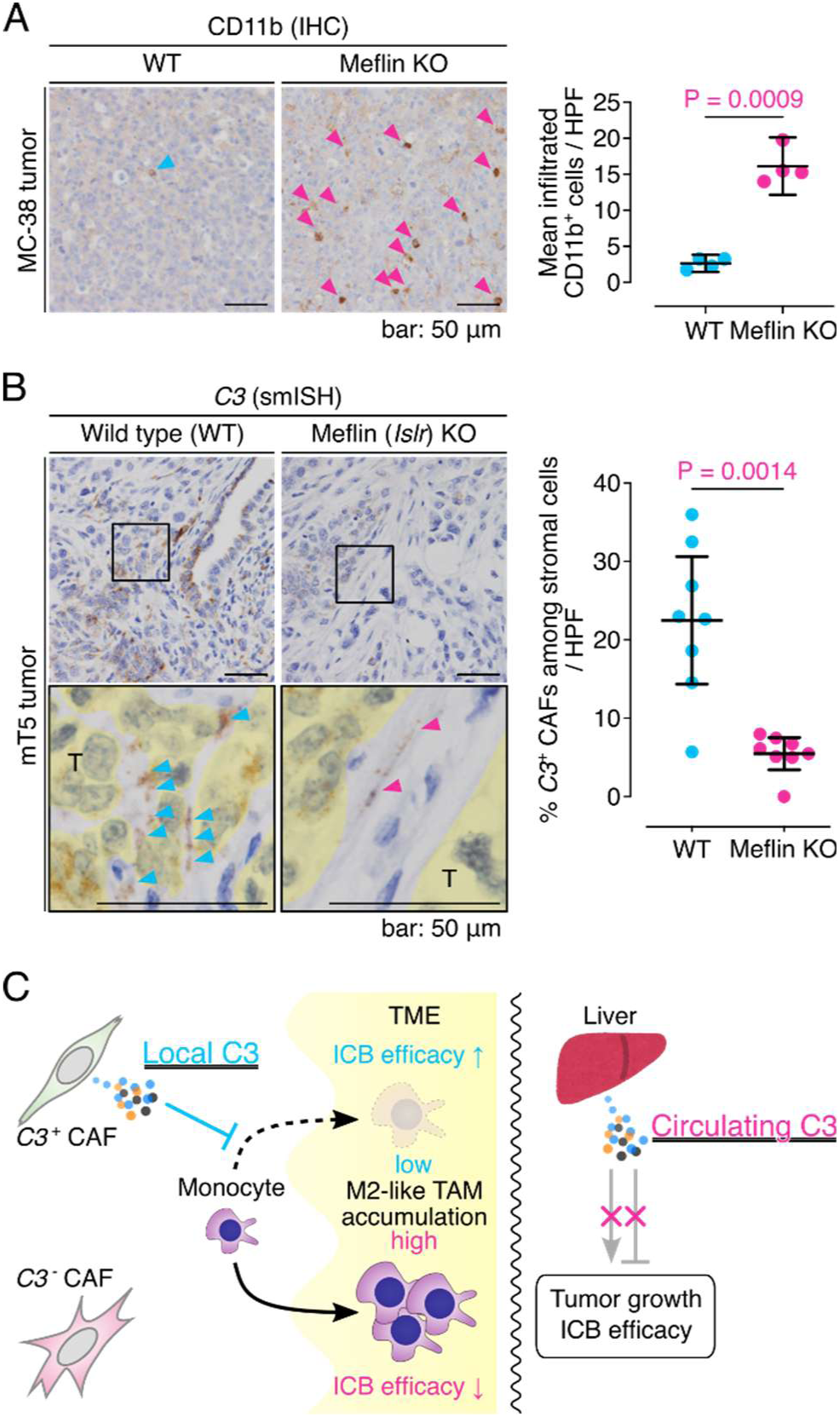
Infiltration of CD11b^+^ cells and *C3*^+^ stromal cells in tumors developed in Meflin KO mice. (A) Tissue sections from MC-38 tumors developed in WT and Meflin KO mice were stained for CD11b using IHC (left), followed by quantification (right) (n = 4 mice, mean of 4 HPFs/mouse). Representative IHC images are shown. CD11b^+^ cells are indicated using arrowheads. (B) Tissue sections from mT5 tumors developed in WT and Meflin KO mice were stained for *C3* using smISH (left), followed by quantification (right). Representative IHC images are shown. Boxed areas are magnified in the lower panels. Arrowheads indicate *C3*^+^ cells. T denotes tumor. (C) Graphical abstract. The data shown in the present study suggested that local, not circulating, complement C3 governs immune checkpoint blockade efficacy. *C3*^+^ cancer-associated fibroblasts (CAFs) inhibit monocyte infiltration into the tumor microenvironment (TME) via the iC3b-CR3 signaling, reducing immunosuppressive M2-like tumor-associated macrophage (TAM) accumulation and enhancing ICB efficacy. In contrast, TME enriched with C3^-^ CAFs shows increased monocyte infiltration, promoting ICB resistance. Notably, hepatocyte-derived circulating C3 does not influence ICB therapy outcomes, highlighting the distinct role of locally produced C3 in cancer immunotherapy.

**Table S1.**
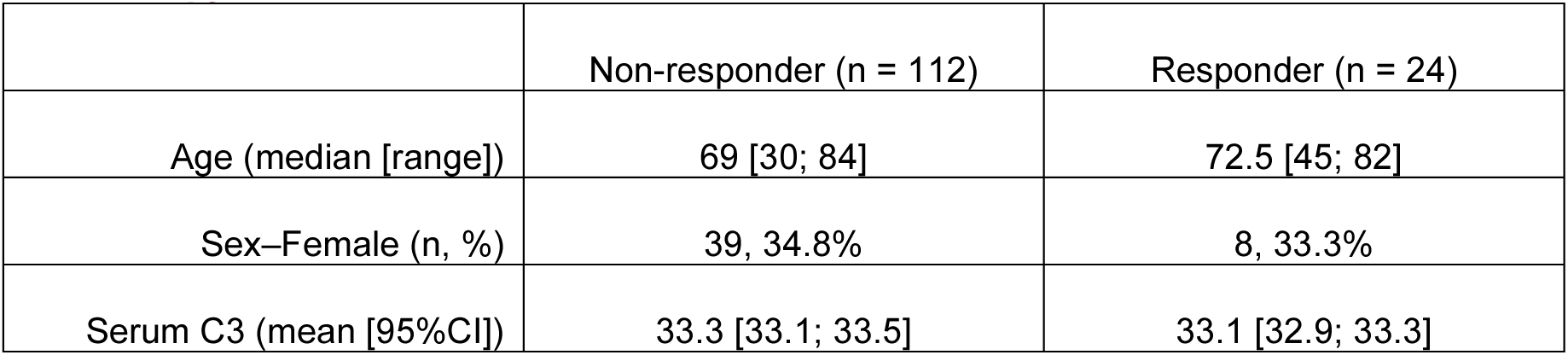
Characteristics of serum cohort of patients with NSCLC who received ICB monotherapy.

|  | Non-responder (n = 112) | Responder (n = 24) |
| --- | --- | --- |
| Age (median [range]) | 69 [30; 84] | 72.5 [45; 82] |
| Sex–Female (n, %) | 39, 34.8% | 8, 33.3% |
| Serum C3 (mean [95%CI]) | 33.3 [33.1; 33.5] | 33.1 [32.9; 33.3] |

**Table S2.** Characteristics of tissue cohort of patients with NSCLC who received ICB monotherapy.

|  | C3 low (n = 14) | C3 high (n = 21) |
| --- | --- | --- |
| Age (median [range]) | 73 [44; 80] | 66 [49; 77] |
| Sex–Female (n, %) | 6, 42.9% | 5, 23.8% |
| Subtype–non-SQ<sup>a</sup> (n, %) | 12, 85.7% | 13, 61.9% |
| BI<sup>b</sup> (median [range]) | 300 [0; 1680] | 900 [0; 2250] |
| OS<sup>c</sup> (median [range]) | 16.5 [0.822; 66.1] | 38.2 [4.96; 81.2] |
| PFS<sup>d</sup> (median [range]) | 1.99 [0.395; 48.8] | 11.4 [0.493; 79.8] |

<sup>a</sup>Non-SQ, non-squamous cell carcinoma.
<sup>b</sup>BI, Brinkman index, defined as (cigarettes per day) × (smoking years).
<sup>c</sup>OS, overall survival (months).
<sup>d</sup>PFS, progression-free survival (months).

## EXPERIMENTAL MODEL AND STUDY PARTICIPANT DETAILS

### Mouse lines

All mice were maintained under specific pathogen-free conditions in the Division of Experimental Animals, Nagoya University Graduate School of Medicine. All experimental protocols were approved by the Animal Care and Use Committee of Nagoya University Graduate School of Medicine (approval number: M240018). Generation of Meflin KO and Meflin (ISLR)-Cre knock-in mice has been previously described^28,51^.

We generated a new conditional C3 knockout (KO) mouse line harboring a C3 floxed allele. Targeted embryonic stem (ES) cell clones (HEPD0812_4_D01, _E04, and _H02) derived from JM8 cells (C57BL/6N background) were purchased from the European Mouse Mutant Cell Repository (EuMMCR). Quality control for correct targeting and homologous recombination in the 5’ and 3’ homology arms was assessed using long-range PCR with EuMMCR. *C3* gene contains 42 exons. EuMMCR designed the targeting construct to insert the Flippase Recognition Target (FRT)-flanked internal ribosome entry site (IRES)-LacZ, neo-resistant genes, and loxP-flanked exons 2–4 between exons 1 and 5. Targeted ES cells (HEPD0812_4_H02) were injected at the Institute of Immunology Co., Ltd. (Tokyo, Japan) into BALB/cA and C57BL/6J blastocysts and chimeric male mice were obtained. The mice were mated with C57BL/6J Rosa26-FLP mice (strain: 009086, JAX, RRID: IMSR_JAX:009086) to excise IRES-LacZ and neo-resistant genes via FRT site recombination. The generated F1 female mice harboring the heterozygous C3 floxed and FLP alleles were crossed with C57BL/6J male wildtype (WT) mice to obtain C57BL/6J mice harboring only the heterozygous C3 floxed allele. After at least three crossings with WT mice, we crossed C57BL/6J *Actb-Cre* transgenic mice (derived from strain: 019099, JAX, RRID: IMSR_JAX:019099), C57BL/6J *Alb1-Cre* transgenic mice (strain: 016832, JAX, RRID: IMSR_JAX:016832), and C57BL/6J Meflin (*Islr*)*-Cre* knock-in mice to obtain systemic C3 KO, hepatocyte-specific C3 KO, and Meflin-lineage C3 KO mice, respectively.

Genomic DNA extracted from mouse tails was used for PCR-based genotyping. The primer sequences used were as follows: WT/floxed C3 forward, 5’-CCAACAACCCTCCAGTCCTC-3’; WT/floxed C3 reverse, 5’-CCAGTGCCAGGTTCAGAGAG-3’; excised C3 forward, 5’-GGCGCATAACGATACCACGA-3’; excised C3 reverse, 5’-CAGAAGTGGGAGCCACAGAG-3’. The PCR product sizes of the WT C3, floxed C3, and excised C3 alleles were 322, 512, and 279 bp, respectively.

### Patient samples

This study was conducted in accordance with the Declaration of Helsinki and approved by the Ethics Committee of Nagoya University Graduate School of Medicine (approval numbers: 2018-0429 for serum analysis and 2017–0127 for tissue analysis). Clinical information and patient samples were collected with permission from the Institutional Review Board-approved protocols in accordance with patient consent. A detailed report on a tissue analysis cohort of patients with NSCLC who received ICB monotherapy (nivolumab, pembrolizumab, or atezolizumab) has been published previously^27^. Serum samples were prospectively collected from patients with NSCLC who received ICB monotherapy at Nagoya University Hospital between November 2015 and September 2020 to clarify the clinical features of endocrine immune-related adverse events (approval number: 2015-0273, UMIN000019024)^52,53^. All patients in this cohort provided written informed consent for serum collection and the secondary use of residual specimens.

## METHOD DETAILS

### Animal experiments and analysis

In vivo tumor studies were performed as follows: 6-week-old female mice were inoculated subcutaneously in their right flanks with 1.0 × 10^6^ MC-38 cells suspended in 100 μL of PBS (day 0). Volumes of the MC-38 tumors were measured and calculated every 3 days from day 4 to 19 using the modified ellipsoid formula 1/2 × (length × width2). Tumor sizes were then measured and calculated every 2 to 3 days. Mice with tumor volumes > 2,000 mm3 were euthanized. Animals whose tumors were ulcerated with bleeding before the volume reached 2,000 mm3 were terminated and included in the survival analysis. Sample sizes were determined based on previous experience with similar experimental models to detect biologically meaningful effects while minimizing the use of experimental animals in accordance with ethical guidelines.

To investigate the efficacy of ICB therapy in the MC-38 subcutaneous tumor model, αPD-1 (clone RMP1-14, 114122, BioLegend, RRID: AB_2800575) and isotype control (clone RTK2758, 400574, BioLegend, RRID: AB_11147167) antibodies were administered intraperitoneally to the mice on days 4, 7, and 10 at a dose of 10 μg/g body weight (BW). C3 inhibitor (AMY-101, CS-0119743, ChemScene) was administered subcutaneously near the tumor site on days 0, 1, 4, 7, 10, and 13 at a dose of 4 μg/gBW (Figure 1), CD11b depletion antibody (clone M1/70, 101272, BioLegend, RRID: AB_2561479) was administered intraperitoneally to mice on days 2, 4, 7, 10, and 13 at a dose of 10 μg/gBW (Figure 7), and CD11b partial agonist (ADH-503 or Leukadherin-1, S8306, Selleck) was administered intraperitoneally to mice every day from day 2 to 13 at a dose of 2 μg/gBW (Figure 7).

A generalized linear mixed-effects model was used to evaluate the effects of temporal changes (time), αPD-1 treatment, and other variables on tumor growth. Log-transformed (tumor volume +1) was used as the dependent variable, with days post-inoculation (*time*), αPD-1 treatment (*P*), other variables (*V*), and their interactions as fixed effects. Individual mouse IDs were included as random intercepts to account for repeated measurements and baseline variations among individual mice. A random intercept structure was chosen over more complex random effect structures (such as random slopes), based on model convergence and parsimony considerations. The model is specified as follows:

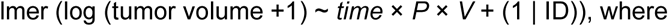

(1 | ID) denotes the random intercept of each mouse. In each experiment, *V* was assigned to each variable: C3 KO status (systemic C3 KO or hepatocyte C3 KO versus control) for Figure 1A–C, local C3 inhibition (AMY-101 versus control) for Fig. 1D–F, genotype (Meflin-lineage C3 KO (mKO) versus control) for Figure 3A–D, and CD11b manipulation (M1/70 or ADH-503 versus PBS) for Figure 7A–D.

As shown in Figure 3A–D, the control mouse data included both blinded and open-label experiments. Because the preliminary analysis showed no significant differences between these conditions (*time* × *control_open*, P = 0.721; *P* × *control_open*, P = 0.846; *time* × *P* × *control_open*, P = 0.372), these data were combined into a single control group for the final analysis to avoid overcomplicating the analysis. Model diagnostics identified statistical outliers that represented the inherent biological variability in treatment responses: complete responses in the mKO group, which showed a lower CR rate than that in the control group, and cases of treatment resistance in the control group, where a favorable therapeutic response was more common. These outliers were retained in the analysis because they represent biologically meaningful variations in treatment response rather than technical artefacts.

All experiments were conducted in multiple independent sessions, each including the WT control group, to ensure reproducibility. Regression lines were used to fit a linear profile to the time course of the logarithm-transformed tumor volumes in each group. All statistical analyses were performed using R (version 4.4.0) with lmerTest package (version 3.1-3)^54^ and lme4 package (version 1.1-35.5)^55^. Statistical significance was set at P < 0.05. Growth curves were visualized using GraphPad Prism 9.

### Cell lines

MC-38 (ENH204, Kerafast, RRID: CVCL_B288), a murine colon cancer cell line; mT5 (generously provided by Drs. David Tuveson (Cold Spring Harbor Laboratory) and Chang-Il Hwang (UC Davis College of Biological Sciences, Davis, CA)), a murine pancreatic cancer cell line; Hep G2 (RBC1886, RIKEN Bioresource Research Center, RRID: CVCL_0027), a human hepatoblastoma cell line, were maintained in DMEM (Nakalai Tesque) supplemented with 10% heat-inactivated fetal bovine serum (FBS) (Gibco). THP-1 (RRID: CVCL_0006, a human monocytic leukemia cell line generously provided by Dr. Fumiya Ito (Nagoya University Graduate School of Medicine)) was maintained in RPMI-1640 medium (Gibco) supplemented with 10% heat-inactivated FBS. All cell lines were routinely screened for mycoplasma contamination through 4, 6-diamidino-2-phenylindole (DAPI) staining.

### Biochemical analysis

Complement C3 concentrations were measured using mouse C3 ELISA kits (ab157711 for plasma and ab263884 for culture supernatant, Abcam) or a human C3 ELISA kit (ab108823, Abcam) according to the manufacturer’s protocols. Blood samples were collected in EDTA-2K tubes and centrifuged at 2,000 g for 10 min at 4 °C. Plasma samples were diluted to 1:50,000 in diluent buffer provided in each kit. Cell culture supernatants were collected after 24 hours of incubation, centrifuged at 2,000 g for 10 min to remove debris, and diluted to fall within the standard curve range. The optical density was measured at 450 nm with path length correction at 977/900 nm using a Cytation 5 microplate reader (BioTek Instruments, Inc., VT, USA). All measurements were performed in triplicate.

Cell culture supernatants were collected after 24 hours of incubation, centrifuged at 2,000 × g for 30 min, and proteins were precipitated using 4 volumes of cold acetone for 60 min at -20 °C. After centrifugation at 15,000 × g for 15 min at 4 °C and air-drying for 10 min, the precipitates were dissolved in 3 × SDS sample buffer (187.5 mM Tris-HCl pH 6.8, 6% SDS, 30% glycerol, and 0.03% bromophenol blue) containing 125 mM dithiothreitol (#7722, Cell Signaling Technology) and heated for 10 min at 70 °C. Proteins were separated on 7.5% sodium dodecyl sulphate-polyacrylamide gel electrophoresis gels and transferred onto nitrocellulose membranes using a wet-transfer system (30 V overnight at 4 °C). The membranes were stained with 0.1% Ponceau S (Sigma-Aldrich) in 1% acetic acid, washed twice with 5% acetic acid, and rinsed twice with water (5 min each). Membranes were blocked with 5% skim milk in TBS containing 0.05% Tween 20 (TBS-T) for 30 min and incubated with an anti-C3 antibody (clone EPR19394, 1:2000, ab200999, Abcam, RRID: AB_2924273) for 60 min at RT. After washing with TBS-T, the membranes were incubated with horseradish peroxidase (HRP)-conjugated swine anti-rabbit antibody (1:2000, P0399, Dako, RRID: AB_2617141) for 30 min at RT. Signals were detected using an ECL reagent (RPN2106, Cytiva) and ImageQuant 800 (Cytiva). Native human C3b (204860-250UG, Sigma-Aldrich) was used as a positive control.

### Single-molecule in situ hybridization and immunohistochemistry

For single-molecule in situ hybridization (smISH), tissue samples were fixed in 10% neutral buffered formalin, embedded in paraffin, and cut to obtain sections with a thickness of 4 μm. The smISH assay was performed using the RNAscope 2.5 HD Detection Reagent-BROWN kit (322310, Advanced Cell Diagnostics) according to the manufacturer’s protocol. For double-staining smFISH, the RNAscope Multiplex Fluorescent v2 Assay Kit (323100, Advanced Cell Diagnostics) was used with Opal 520 (FP1487001KT, Akoya) and Opal 690 (FP1497001KT, Akoya) fluorophores. Probes targeting human *ISLR* (455481 or 455481-C2, Advanced Cell Diagnostics), *C3* (430701, Advanced Cell Diagnostics), mouse *Islr* (450041 or 453321-C2, Advanced Cell Diagnostics), *C3* (417841, Advanced Cell Diagnostics), and *C3* [exon2–4] (1116031-C1, Advanced Cell Diagnostics, newly designed) were used.

For immunohistochemistry (IHC), formalin-fixed paraffin-embedded sections were deparaffinized, rehydrated, blocked with endogenous peroxidase, and subjected to heat-induced epitope retrieval in citrate buffer (pH 6.0) in a hotpot. After blocking with 2.5% serum from the same host as the secondary antibody for 30 min, the sections were incubated with primary antibodies against CD11b (clone EP45, 1:200 dilution, AC-0043RUO, EPITOMICS, RRID: AB_10999891) or CD206/MRC1

(clone E2L9N, 1:400 dilution, 91992, Cell Signaling Technology, RRID: AB_2800175) for 60 min at room temperature. After washing, the sections were incubated with an HRP-conjugated secondary antibody (MP-7451, ImmPRESS HRP Goat Anti-Rabbit IgG Polymer Kit, Vector Laboratories) for 30 min at room temperature. The signals were visualized using a 3,3’-diaminobenzidine (DAB) substrate (SK-4103, ImmPACT DAB EqV, Vector Laboratories). For double IHC, the sections were incubated with antibodies against CD206/MRC1 and CD68 (clone KP1, 1:500 dilution, M0814, Dako, RRID: AB_2314148). After washing, the sections were incubated with HRP- and alkaline phosphatase (AP)-conjugated secondary antibodies (MP-7402 and MP-5401, ImmPRESS HRP Horse Anti-Mouse and AP Horse Anti-Rabbit IgG Polymer Detection Kit, Vector Laboratories) for 30 min at room temperature. The signals were visualized using DAB substrate and ImmPACT Vector Red AP substrate (SK-5105, Vector Laboratories).

For smISH or smFISH quantification, cells harboring more than four dots or clustered signals were considered signal-positive cells. For correlation analysis between *ISLR* and *C3* expression, the proportions of *ISLR*^+^ and *C3*^+^ stromal cells were calculated for each high-power field (x400), excluding morphologically distinct immune, vascular endothelial, and epithelial cells. Pearson’s correlation coefficients were calculated for each cancer type, with 95% confidence intervals. Correlation analysis etween the proportion of *C3*^+^ CAFs and MRC1^+^ M2-like TAM infiltration (determined by IHC) was performed for each cancer type using Spearman’s rank correlation coefficient with 95% confidence. A meta-analysis of the correlations across cancer types was performed using a random-effects model. Forest plots were generated to visualize the correlation coefficients for each cancer type along with the pooled estimates. Statistical analyses were performed using R (version 4.4.0) with meta package (version 7.0.0)^56^. Heterogeneity between cancer types was assessed using the I^2^ statistic and τ^2^ value.

### Single-cell RNA sequencing analysis

We analyzed publicly available scRNA-seq datasets for colorectal cancer (GSE132465)^57^, non-small cell lung cancer (NSCLC, GSE131907)^58^, and pancreatic ductal adenocarcinoma (PDAC, GSE155698)^59^. After removing both low-quality cells and malignant cells, 95,780 stromal cells were obtained. Gene expression data were log-normalized and batch-corrected using R (version 4.4.1) with Seurat package (version 5.1.0)^60^ employing the IntegrateLayers function using the canonical correlation analysis integration method. Cell types were annotated based on canonical marker genes. *ISLR* and *C3* expression patterns were visualized using UMAP dimensionality reduction, and the expression levels of *ISLR* and *C3* across cell types were displayed using dot plots.

For the correlation analysis, we first calculated the mean expression of *C3* in CAFs for each patient. The proportions of M2-like TAMs and total myeloid cells among stromal cells were also calculated for each patient. Myeloid cells and M2-like TAMs were identified based on the expression of *ITGAM* and *ITGAM* with *MRC1*. Pearson’s correlation coefficients between the mean CAF *C3* expression and the proportions of these immune cell populations were calculated for each cancer type. Meta-analyses of these correlations were performed using the random-effects models, and the results were visualized using forest plots with 95% confidence intervals. Statistical analyses were performed using R (version 4.4.0) with meta package (version 7.0.0)^56^. Heterogeneity between cancer types was assessed using the I^2^ statistic and τ^2^ value.

For the analysis of CAF subsets, we used the CELL×GENE Discover Census database (Chan Zuckerberg Initiative, version 2023-12-15)^61^. We extracted annotated fibroblasts from patients with cancer, including those with breast cancer (n = 1)^62^, clear cell renal cell carcinoma (n = 2)^63,64^, non-small cell lung carcinoma (n = 3)^65–67^, and small cell lung cancer (n = 1)^68^. After quality control and normalization, 73,500 cells were obtained. The batch effects between the datasets were corrected using harmony integration, and clusters were identified using the Leiden algorithm.

For each CAF subset, the top 200 differentially expressed genes were identified. Expression values were z-scores normalized across clusters, and the average expression patterns were visualized using a heatmap. The relative expression levels of marker genes were used to annotate the CAF subsets based on previously reported signatures. CAF subsets were annotated based on their expression profiles as: iCAF (*IL1R1*, *IL6*-expressing inflammatory CAFs), sslCAF*^PI16^* (*C3*, *DPT*, *ISLR*, *PI16*-expressing steady state-like CAFs), myCAF (*POSTN*, *ACTA2*-expressing myofibroblastic CAFs), sslCAF*^COL15A1^* (*APOD*, *COL15A1*-expressing steady state-like CAFs), apCAF (*CD74*, *HLA-DRB1*-expressing antigen-presenting CAFs), and vCAF (*MCAM*, *RGS5*-expressing vascular CAFs or CAF-S4).

Co-expression analysis of *C3* with *ISLR* or *DPT* was performed within the CAF population from the CRC, NSCLC, and PDAC datasets, and the sslCAF*^PI16^* subset from the CELL×GENE Discover Census database. Cells were classified as positive for each gene when normalized expression exceeded 1, and the percentage of cells in each co-expression category was calculated relative to the total number of cells in the subset.

Analysis of the CELL×GENE Discover Census database was performed using Python (version 3.10.11) with Scanpy (version 1.9.3), harmony-py (version 0.1.8), SciPy (version 1.13.1), NumPy (version 1.24.3), pandas (version 2.2.2), and anndata (version 0.9.1) packages. Visualizations were generated using Matplotlib (version 3.7.1) and seaborn (version 0.12.2).

### Bulk RNA-seq analysis

Total RNA was extracted from MC-38 tumors that developed in control and mKO mice (n = 2 per group) using TRI-Reagent (Molecular Research Center) following the manufacturer’s protocol. RNA quality was assessed using an Agilent 2100 Bioanalyzer, and samples with an RNA integrity number (RIN) > 6 were used for library preparation. RNA-seq libraries were prepared using the NEBNext Poly(A) mRNA Magnetic Isolation Module (E7490) and NEBNext Ultra II Directional RNA Library Prep Kit (E7760), according to the manufacturer’s instructions. The libraries were sequenced on an Illumina NovaSeq 6000 platform with paired-end 150 bp reads to obtain 6 G bases (40 M reads) per sample.

Raw sequencing reads were quality-controlled using FastQC (version 0.11.7) and trimmed using Trimmomatic (version 0.38) software. The clean reads were mapped to the mouse reference genome (mm10) using HISAT2 (version 2.1.0). Gene-level counts were generated using FeatureCounts (version 1.6.3). Differential expression analysis was performed using R (version 4.2.3) with TCC package (version 1.38.0)^69^, employing a trimmed mean of M-value normalization. The empirical Bayesian method implemented in edgeR (version 3.40.2) was used for differential expression testing, with three iterations of the DEG elimination strategy. Genes with false discovery rate (FDR) values < 0.05 were considered differentially expressed. Gene Ontology over-representative analysis of the upregulated genes was performed using clusterProfiler package (version 4.6.2)^70^. Biological process terms with adjusted P value < 0.01 and Q value < 0.05 were considered significantly enriched. The universe of background genes was set for all genes detected in the RNA-seq analysis.

### CITE-seq analysis of tumor-infiltrating immune cells

MC-38 tumors were mechanically minced and enzymatically digested using the tumor dissociation kit (130-096-730, Miltenyi Biotec) in gentleMACS C-Tubes (130-093-237, Miltenyi Biotec) according to the manufacturer’s instructions with the pre-installed program “37C_h_TDK_4”. Single-cell suspensions were filtered through 70 μm cell strainers and treated with ACK buffer for red blood cell lysis. Dead cells were removed using dead cell removal microbeads according to the manufacturer’s protocol. Following Fc receptor blocking with TruStain FcX PLUS (S17011, #156603, BioLegend, RRID: AB_2783137), cells were stained with TotalSeq antibodies (list shown below, all from BioLegend) for protein detection, and CD45^+^ immune cells were enriched using CD45 MicroBeads and MS columns (130-042-201, Miltenyi Biotec). Cell viability (> 80%) was assessed using Countess before loading onto the 10x Genomics Chromium Next GEM Single Cell 3’ v3.1 platform.

| Antibody | clone | Cat # | RRID |
| --- | --- | --- | --- |
| TotalSeq-B0001 anti-mouse CD4 Antibody | RM4-5 | 100573 | AB_2813914 |
| TotalSeq-B0002 anti-mouse CD8a Antibody | 53-6.7 | 100783 | AB_2832269 |
| TotalSeq-B0013 anti-mouse Ly-6C Antibody | HK1.4 | 128053 | AB_2832462 |
| TotalSeq-B0014 anti-mouse/human CD11b Antibody | M1/70 | 101273 | AB_2819781 |
| TotalSeq-B0015 anti-mouse Ly-6G Antibody | 1A8 | 127659 | AB_2819864 |
| TotalSeq-B0103 anti-mouse/human CD45R/B220 Antibody | RA3-6B2 | 103271 | AB_2832306 |
| TotalSeq-B0106 anti-mouse CD11c Antibody | N418 | 117359 | AB_2813987 |
| TotalSeq-B0118 anti-mouse NK-1.1 Antibody | PK136 | 108763 | AB_2819814 |
| TotalSeq-B0182 anti-mouse CD3 Antibody | 17A2 | 100257 | AB_2813912 |
| TotalSeq-B0184 anti-mouse CD335 (Nkp46) Antibody | 29A1.4 | 137641 | AB_2860685 |
| TotalSeq-B0091 Mouse IgG2a, κ isotype Ctrl | MOPC-173 | 400291 | AB_3097066 |
| TotalSeq-B0095 Rat IgG2b, κ Isotype Ctrl | RTK4530 | 400689 | AB_3097109 |
| TotalSeq-B0238 Rat IgG2a, κ Isotype Ctrl | RTK2758 | 400581 | AB_3097104 |
| TotalSeq-B0241 Armenian Hamster IgG Isotype Ctrl | HTK888 | 400979 | AB_3097125 |

Gene expression and antibody-derived tag (ADT) libraries were prepared using the Chromium Next GEM Single Cell 3’ Reagent Kits v3.1 (Dual Index) with Feature Barcode technology for Cell Surface Protein (10x Genomics, CA, USA) according to the manufacturer’s instructions. Libraries were sequenced on an Illumina NovaSeq 6000 platform with read lengths of R1(28bp)-I1(8bp)-I2(8bp)-R2(90bp). Raw base call files were processed using Cell Ranger (version 7.0.1) and bcl2fastq (version 2.20.0). Quality control metrics showed > 92% of bases with Q30 quality scores for gene expression libraries and > 86% for ADT libraries.

Single-cell gene expression and protein data were analyzed using R (version 4.4.2) and Python (version 3.10.11). After quality control filtering, including the removal of cells with high mitochondrial gene content (>7.5%), abnormal RNA/protein count distributions (>5 median absolute deviations), and potential doublets identified using scDblFinder (version 1.14.0)^71^ and SingleCellExperiment (version 1.22.0)^72^, 12,528 cells from four individual tumors (n = 2 per genotype group) were obtained. Ambient RNA contamination was corrected using SoupX (version 1.6.2)^73^. Integrated analysis of RNA and protein expression was performed using totalVI implemented in scvi-tools (version 1.1.2)^74^. The model employs a hierarchical Bayesian model with default priors, including log-normal priors for scaling factors and normal priors for latent variables. Following model training and convergence, a low-dimensional latent representation was extracted. Genotype-corrected normalized expression values for both RNA and protein modalities were obtained by averaging 25 samples from the posterior distribution. Cell clustering was performed using the Leiden algorithm in the totalVI latent space. Major immune cell populations were identified based on their transcriptional profiles (using differentially expressed genes with a threshold of delta=0.5), and surface protein expression patterns were visualized using dot and matrix plots.

Cell type composition analysis between genotypes was conducted using scCODA (version 0.1.9)^75^, with Th17 cells selected as the automatic reference cell type. The analysis employed default priors, and Markov chain Monte Carlo sampling was performed using the No-U-Turn Sampler (NUTS) with 11,000 iterations. Credible effects were determined at a false discovery rate of 0.05. Cell-cell interaction analysis was performed using CellPhoneDB implemented in liana (version 1.2.1)^76^ to identify significant receptor-ligand interactions (P < 0.05), followed by network analysis using NetworkX (version 3.1)^77^. The PageRank algorithm was used to identify crucial cellular hubs in the tumor microenvironment, with interaction strengths weighted by the mean expression of significant ligand-receptor pairs.

Bayesian analyses (totalVI and scCODA) were performed with GPU acceleration using PyTorch (version 2.2.1) and TensorFlow (version 2.12.0) backends, respectively. The analysis pipeline was implemented using JupyterLab (version 4.1.6) with rpy2 (version 3.5.11) and anndata2ri (version 1.1) for R-Python integration. Data processing and visualization were performed using Python-based tools, including Scanpy (version 1.9.3), muon (version 0.1.3), and anndata (version 0.9.1). Data manipulation and visualization were conducted using NumPy (version 1.24.3), pandas (version 2.2.2), Matplotlib (version 3.7.1), and seaborn (version 0.12.2).

### Flow cytometry analysis

To isolate cells from tumors for FCM analysis, tumors were mechanically minced and enzymatically digested using the tumor dissociation kit (130-096-730, Miltenyi Biotec) in gentleMACS C-Tubes (130-093-237, Miltenyi Biotec) according to the manufacturer’s instructions using the pre-installed program “37C_h_TDK_4”. Single-cell suspensions were filtered through 70 μm cell strainers and treated with ACK buffer for red blood cell lysis.

FCM staining and analysis were performed using conventional procedures. The cells were washed with FACS buffer (0.5% BSA and 2 mM EDTA in PBS) and blocked with TruStain FcX PLUS (S17011, #156603, BioLegend, RRID: AB_2783137). Cells were then stained with cell-surface antibodies and the fixable viability dye eFlour 780 (65-0865-14, eBioscience). For intracellular staining of iNOS, cells were fixed and permeabilized using the Cytofix/Cytoperm Fixation/Permeabilization Kit (554714, BD, RRID: AB_2869008) according to the manufacturer’s instructions, followed by staining with monoclonal antibodies against iNOS. After washing, the cells were analysed using a BD LSR Fortessa X-20 flow cytometer (BD Biosciences) with FACSDiva (version 9.0) and FlowJo (version 10.9.0) software. In this study, the following anti-mouse antibodies labelled with fluorescent dyes were used (all at 1:100 dilution, unless otherwise specified): CD45–PerCP/Cyanine5.5 (clone 30-F11, 103132, RRID: AB_893340), F4/80–APC (clone BM8, 123116, RRID: AB_893481), CD11b–Brilliant Violet 421 (clone M1/70, 1:200 dilution, 101251, RRID: AB_10897942), CD206–PE (clone C068C2, 141705, RRID: AB_10895754) (all from BioLegend); iNOS–Alexa Fluor 488 (clone CXNFT, 53-5920-80, eBioscience, RRID: AB_2574422).

### Clinical data analysis

Overall survival (OS) was defined as the time from ICB therapy initiation to death from any cause, whereas progression-free survival (PFS) was defined as the time from ICB therapy initiation to documented disease progression or death from any cause. Patients who survived without documented disease progression were censored at the last follow-up visit. Treatment responses were evaluated by the attending physicians based on clinical assessments, including radiological findings. Responders were defined as patients who achieved complete response (CR) or partial response (PR), whereas non-responders were defined as those who showed stable disease (SD) or progressive disease (PD). Multivariate Cox proportional hazard models were used to assess the associations of serum C3 levels with OS and PFS after adjusting for sex and age. OS, PFS, and response rates were compared between patients with high and low proportions of *C3*^+^ CAFs in the stroma.

For survival analysis using the Kaplan-Meier Plotter database^78^, we analyzed 933 patients with solid tumors who received immunotherapy and had evaluable C3 expression data. Based on stromal C3 expression normalized to the expression of the fibroblast marker COL1A1, these patients were classified into the C3 high (n = 625) and C3 low (n = 308) groups. The 2:1 ratio was selected to match the distribution observed in the NSCLC tissue cohort.

### Mass spectrometric quantification of serum C3

Ten microliters of serum sample were treated using a high-select top-14 abundant protein depletion column (A36370, Thermo Fisher Scientific) and concentrated through ultrafiltration. The total protein content was quantified using the Bradford method. After incubating 12.5 μg of protein in denaturing buffer (25 mM NH4HCO3, 8 M Urea, and 5 mM DTT) at 37 °C for 1 hour, iodoacetic acid was added to achieve a final concentration of 20 mM, and the mixture was incubated in the dark at room temperature for 30 min. The mixture was then diluted four times with 50 mM NH4HCO3, followed by the addition of 0.5 μg of Trypsin Gold (V5280, Promega), and incubated overnight at 37 °C. The reaction was quenched with trifluoroacetic acid to a final concentration of 0.17% (v/v). Five pmol of Peptide Retention Time Calibration Mixture (88321, Thermo Fisher Scientific) was added, and the mixture was concentrated by evaporation. The samples were purified using Pierce C-18 spin columns (89873, Thermo Fisher Scientific). After drying by evaporation, the pellet was dissolved in 25 μL of 0.1% formic acid.

Liquid chromatography-mass spectrometry (LC-MS) was performed using an EASY-nLC 1200, Orbitrap Eclipse Tribrid Mass Spectrometer, and a FAIMS Pro interface (Thermo Fisher Scientific). One microliter of each sample was injected and separated using an Acclaim PepMap C18 column (75 μm × 20 mm, 3 μm, 164946, Thermo Fisher Scientific) as the trap column and a NANO HPLC Capillary Column (75 μm × 125 mm, 3 μm, NTCC-360/75-3-125, Nikkyo Technos) as the analytical column at 35 °C. The mobile phases were 0.1% (v/v) formic acid in water (mobile phase A) and 0.1% (v/v) formic acid in 80% acetonitrile (mobile phase B). The gradient condition was as follows: mobile phase B of 6% (0–1 min)–31% (91 min)–50% (91.1–96.1 min)–90% (96.1–115 min). The flow rate was set at 300 nL/min. The MS condition was in the positive mode with a spray voltage of 2000 V, and the ion transfer tube was set to 275 °C. For MS1 acquisition, spectra were collected using an orbitrap analyzer in the range 375–1500 m/z at a resolution of 120,000, and AGC target set to 4e5. For MS2 acquisition, spectra were collected using an ion trap analyzer at a turbo scan rate with a collision energy of 35% and AGC target set to 1e4. The dynamic exclusion time was set to 20 s, and the FAIMS CVs were 40, 60, and 80 V. Collected data were processed using Proteome Discoverer 2.4 (Thermo Fisher Scientific). Proteins were searched against UniProt Swiss-Prot human proteome (version Oct. 25, 2017). Relative protein abundances were calculated based on the intensities of identified peptides and normalized to the internal standard peptides (Peptide Retention Time Calibration Mixture). These normalized values were then log2-transformed for statistical analysis. Proteins with a missing rate >30% were excluded; C3 had a complete detection rate (0% missing). The remaining missing values were imputed using missForest package (version 1.5)^79^ in R (version 3.6.0). The batch effect was corrected using ComBat function from sva package (version 3.32.1)^80^.

Serum C3 levels were extracted from the proteomic data of each sample. Based on clinical information, the patients were classified as responders or non-responders. Two one-sided tests (TOST) were performed using R (version 4.4.0) with TOSTER package (version 0.8.4)^81^ employing equivalence bounds set at ± 3%, which is less than one-fifth of the coefficient of variation calculated from the reference range (73–138 mg/dL) of healthy Japanese individuals. Graph visualization was performed using GraphPad Prism 9.

### TCGA database analysis

The Cancer Genome Atlas (TCGA) datasets were accessed through the TCGAbiolinks package (version 2.26.0)^82^. We obtained RNA-seq data (HTSeq-FPKM) for each cancer type and transformed them into transcripts per million. To normalize stromal gene expression, *C3* expression values were divided by COL1A1 expression values to account for variations in stromal content between samples. Similarly, MRC1 expression was normalized to PTPRC (CD45) expression to account for differences in immune cell infiltration. Samples with zero or missing values for any of these genes were excluded from the analysis. Spearman’s rank correlation coefficients were calculated for the correlation analysis between normalized C3 and MRC1 expression across different cancer types.

Based on the biological context of our study, we excluded sarcomas (due to the potential expression of CAF-like genes by tumor cells), brain tumors (where TAMs are predominantly derived from tissue-resident microglia), and hematological malignancies (which lack or contain minimal CAF populations). A meta-analysis of the correlation coefficients was performed using a random-effects model. Forest plots were generated to visualize the correlation coefficients and their 95% confidence intervals for each cancer type, along with the pooled estimates. Statistical analyses were performed using R (version 4.4.0) with meta package (version 7.0.0)^56^. Heterogeneity between cancer types was assessed using the I^2^ statistic and τ^2^ value.

### Primary cell preparation

Primary peritoneal CD11b^+^ cells were prepared using thioglycolate broth (#5601, Nissui Pharmaceutical). Briefly, 2 ml of 3% thioglycolate broth was injected into the peritoneal cavity of 7-week-old female C57BL/6J WT mice. Approximately 36 hours later, we sacrificed the mice and injected 5 ml ice-cold PBS with 2 mM ethylenediaminetetraacetic acid (EDTA) buffer into the peritoneal cavity, gently massaged the abdomen, and collected the liquid containing the peritoneal inflammatory cells. Cells were centrifuged (300 × g, 5 min, 4 °C) and resuspended in ammonium-chloride-potassium lysis buffer to break down the red blood cells. The cells were centrifuged (300 × g, 5 min, 4 °C) again and resuspended in ice-cold PBS with 2 mM EDTA buffer containing CD11b microbeads (130-049-601, Miltenyi Biotec). We then isolated peritoneal CD11b+ cells using MACS MS columns according to the manufacturer’s instructions and incubated them in a plastic dish for 1 hour in RPMI-1640 supplemented with 10% heat-inactivated FBS to remove firmly adherent cells. Non-attached cells were collected and subjected to a transwell migration assay.

Primary human CD14^+^ monocytes were purchased from Lonza (2W-400B, batch numbers 19TL113254 and 22TL108493). The cells were thawed, incubated in plastic dishes for 24 hours, and subjected to transwell migration assays.

### RNA interference

For C3 knockdown, Hep G2 cells were transfected with control siRNA (1027310) or *C3*-specific siRNA (SI02777075 and SI03076689) purchased from QIAGEN (FlexiTube siRNA, Cat. no. 1027417) using Lipofectamine RNAiMAX Transfection Reagent (13778-150, Thermo Fisher Scientific) according to the manufacturer’s instructions. Knockdown efficiency was confirmed using ELISA and western blotting. For a stable CD11b knockdown, THP-1 cells were transduced with lentiviral particles containing either scrambled short hairpin RNA (shRNA) or CD11b (ITGAM)-targeted shRNA. All lentiviral vectors containing shRNA target sequences were designed and synthesized by VectorBuilder. The target sequences for shRNA were as follows: sh_scramble, 5’-CCTAAGGTTAAGTCGCCCTCG-3’; sh_ITGAM #1, 5’-CAACTGTGATGGAGCAATTAA-3’; sh_ITGAM #2, 5’-AGCGCTGCCATCACCTCTAAT-3’; sh_ITGAM #3, 5’-GAAACTACAGTTGCCGAATTG-3’. Lentiviral vectors were transfected into 293FT cells (R70007, Thermo Fisher Scientific, RRID: CVCL_6911) with ViraPower Lentiviral Packaging Mix (JPG0035, Thermo Fisher Scientific) using Lipofectamine 2000 (11668019, Thermo Fisher Scientific). Knockdown efficiency was confirmed using flow cytometry.

### Cell assays

Directional cell migration was assessed using transwells with 5.0 μm pore size inserts (Corning). The cells and media used in the experiments are indicated in each panel of Figures 5,6. After 3 hours of incubation, the transmigrated cells were fixed, stained with hematoxylin, and counted in 4–6 random fields per well under a microscope.

Random cell migration was monitored using the Live-Cell Imaging and Analysis Instrument Incucyte SX5 (Sartorius, Göttingen, Germany). THP-1 cells were cultured in 1% C3-depleted serum-containing media with or without iC3b (2 µg/mL, A115, CompTech) and/or BAY 61-3606 (1 μM, S7006, Selleck) pretreatment. Time-lapse images were acquired every 2 min for 2 hours with two fields of view per well and two wells per condition. Cell trajectories were manually tracked using the Fiji/ImageJ software (2.14.0/1.54f) by randomly selecting eight cells per field (32 cells per condition). The obtained coordinate data were processed using R (version 4.4.0) with dplyr package (version 1.1.4) and ggplot2 package (version 3.5.1) to calculate and visualize individual cell trajectories and total migration distances.

For adhesion assays, THP-1 cells were seeded in 1% C3-depleted serum-containing media with or without iC3b and/or BAY 61-3606 (1 μM) pretreatment. After incubation for 1 hour, bright-field images were immediately captured using a BZ-X710 microscope (KEYENCE, Osaka, Japan) with oblique illumination. The cells were then gently washed with PBS to remove non-adherent cells, and the same fields were imaged again. The adherent cells were counted in three low-power fields per well.

## QUANTIFICATION AND STATISTICAL ANALYSIS

GraphPad Prism 9 was used for statistical analysis unless otherwise specified. The data are expressed as means with 95% confidence intervals (CIs), medians with ranges, or numbers with percentages, as appropriate. Comparisons between groups were performed using a two-tailed unpaired t-test with Welch’s correction unless otherwise specified. Survival was analyzed using the Kaplan-Meier approach and the log-rank Mantel-Cox test. For multiple testing, the Holm–Bonferroni method was employed. Effect sizes, including hazard ratios, odds ratios, and correlation coefficients, are reported with their 95% CIs. Detailed statistical methods for specific analyses are described in the respective method sections. Statistical significance was set at P < 0.05.

